# Modelling human and non-human animal network data in R using STRAND

**DOI:** 10.1101/2022.05.13.491798

**Authors:** Cody T. Ross, Richard McElreath, Daniel Redhead

**Affiliations:** Department of Human Behavior, Ecology, and Culture, Max Planck Institute for Evolutionary Anthropology. Leipzig, Germany

**Keywords:** social networks, animal networks, social relations, R software, generative models

## Abstract

1. There have been recent calls for wider application of generative modelling approaches in applied social network analysis. These calls have been motivated by the limitations of contemporary empirical frameworks, which have generally relied on *post hoc* permutation methods that do not actively account for interdependence in network data. At present, however, it remains difficult for typical end-users—e.g., field researchers—to apply generative network models, as there is a dearth of openly available software packages that make application of such methods as simple as other, permutation-based methods.
2. Here, we outline the `STRAND` R package, which provides a suite of generative models for Bayesian analysis of human and non-human animal social network data that can be implemented using simple, base `R` syntax.
3. To facilitate ease-of-use, we provide a tutorial demonstrating how `STRAND` can be used to model binary, count, or proportion data using stochastic blockmodels, social relations models, or a combination of the two modelling frameworks.

## Introduction

The application of theory and methods from network science (i.e., social network analysis) to ethological data has led to important advances in our understanding the structural features of animal societies. Similarly, the role that sociality— broadly conceived—plays in the differential success and survival of individuals and groups (e.g., Clutton-Brock 2009) has been a topic of perennial interest across the social, behavioural and biological sciences. By quantifying the social interactions (i.e., ties or edges) that are observed between individuals (i.e., nodes or vertices), researchers can more formally study how various dyadic phenomena are related to one another (e.g., Smith-Aguilar et al. 2019) and to key individual-level properties (e.g., Pike et al. 2008).

Recent network-based research has, for example, advanced theory on how social relationships—both positive (e.g., food sharing and grooming) and negative (e.g., agonistic behaviours)—guide the emergence and maintenance of social hierarchies (Kawakatsu et al. 2021; Redhead and Power 2022), influence the spread of disease (Read et al. 2008; Silk et al. 2017) and adaptive information (Waters and Fewell 2012; Hobaiter et al. 2014), and explain how individual actions culminate in group-wide movement patterns (Jacoby and Freeman 2016; Strandburg-Peshkin et al. 2015). To address these topics, and many others, network data are rapidly being compiled across a broad range of taxa (Sah et al. 2019). Given the flexibility of network-based frameworks for understanding behaviour, social network analysis has become one of the most popular areas of research in animal behaviour, behavioural ecology, and the quantitative evolutionary and social sciences more broadly.

### Inferential concerns and topics of debate

While network analytical tools have great potential for advancing our understanding social behaviour across taxa, there are many statistical complexities inherent in such approaches. Network data are highly interdependent, and cannot be modelled using standard statistical approaches that assume uncorrelated residuals. Standard practices within the field often ignore such issues. For example, it is common for researchers to regress outgoing ties on incoming ties to estimate reciprocity (e.g., Carter and Wilkinson 2013), but such regressions are known to suffer from residual confounding (see Koster and Leckie 2014). Similarly, it is common for researchers to correct for sampling effort by creating a Simple Ratio Index (SRI; Cairns and Schwager 1987; Whitehead and James 2015; Farine and Whitehead 2015), but such indices are well-know to divide out sample size and give the weakest data-points disproportionate weight in downstream analyses (Hart et al. 2021b).

Permutation methods—such as the quadratic assignment procedure (QAP; Hubert and Schultz 1976; Krackardt 1987; Dekker et al. 2003)—have been used to “account for” the typical non-independence of network data (see Farine and Carter 2022; Farine and Whitehead 2015; Sosa et al. 2021). However, such *post hoc* methods do not actually permit unbiased estimation of generative model parameters (see Hart et al. 2021a), a point raised by a founder of the method thirty years ago (Krackhardt 1992). While there remains much lingering debate about the usefulness of permutation methods in applied network analysis, it is widely agreed upon that empirical data of all forms are best analysed using scientifically informed generative models (e.g., Reilly and Zeringue 2005). In the case of social network data, such models should *actively* account for non-independence of data points via correlated random effects at both the node and dyad level during the process of model fitting (see McElreath 2020, for a textbook example).

Permutation methods rose to prominence in large part for historical reasons, as correlated random-effects models were difficult to fit using early computer software. In fact, nuanced generative models of social networks have been available for decades, but Bayesian estimation of these models remained computationally intractable until only recently. As such, there is still a dearth of freely-available open-source software for implementing such generative modelling approaches as data analysis tools. To resolve this issue, we draw upon classic generative modelling approaches for network data, and integrate them with contemporary tools for Bayesian model fitting (Stan Development Team 2021b). In doing this, we have developed an R package, `STRAND`, that allows end-users to build complex network analysis models using simple `lm`-style syntax in base R, simplifying the process of modelling empirical network data.

### Generative modelling approaches

There are a plethora of generative network models that reflect complex data generating procedures (see Newman 2018). Through Bayesian inversion (Allmaras et al. 2013), these models can be used as analytic tools that support statistical inference on the basis of empirical data. One such model that holds particular promise for research on animal social networks is the *social relations model* (Kenny and La Voie 1984; Snijders and Kenny 1999), which examines and accounts for correlations in node-level and dyad-level random effects. Across many contexts, animal social networks may further be partitioned into observable subgroups—such as coalitions (Kajokaite et al. 2019), matrilines (Ilany et al. 2021), and groupings based on identity or physical location (De Dreu and Triki 2022). These groupings may create gross community structure in networks, whereby individuals preferentially interact with those in their own sub-groups (to a greater or lesser extent across contexts and relationship types; e.g., Pisor and Ross 2021). Given this, *stochastic blockmodels* show further promise for animal social network analysis (Pearl and Schulman 1983), as they capture these higher order structures (Peixoto 2019). Together, these approaches provide a framework for a more direct analysis of the mechanisms involved in the data generating process of social relationships.

### Our contribution

In order to address the concerns outlined above, and to make data analysis using complex network models easier for end-users, we have introduced an R package for Bayesian social network analysis, `STRAND`, that facilitates applications of generative network modelling approaches. The `STRAND` package supports stochastic blockmodelling, social relations modelling, and more complex latent-network modelling approaches (see Redhead et al. 2021).

In this paper, we outline how stochastic blockmodels and social relations models can be fit to both human and non-human animal network data to answer key research questions. By presenting a tutorial for fitting these models to each of the three most commonly collected types of outcome data used in studies of animal behaviour and behavioural ecology—i.e., binary tie data (e.g., via self-reports), count data (e.g., through behavioural observations over a standardised time window), and proportion data (e.g., counts of behavioural observations where the sampling rate is variable across possible dyads)—we hope to inspire more wide-spread application of principled generative network models in empirical research.

Our approach to network modeling here complements that of the `BISoN` team (Hart et al. 2021b), who have also developed a suite of network analysis models using Bayesian methods. Our R package provides additional functionality to typical end-users, however, in that it integrates new Bayesian network analysis models on the ‘back-end’ with a user-friendly ‘front-end’ interface, that allows even casual R users to specify complex network models (which may include a variety of block-, individual-, and dyad-level covariate data) on-the-fly, using nothing more than base R syntax.

## Using STRAND

Much of the functionality of `STRAND` is made possible by `Stan` (Stan Development Team 2021b) and `CmdStanR` (Stan Development Team 2021a). Users must install these programs prior to installing `STRAND`. Installation and loading of `STRAND` is then simple: just run three lines of code from R:

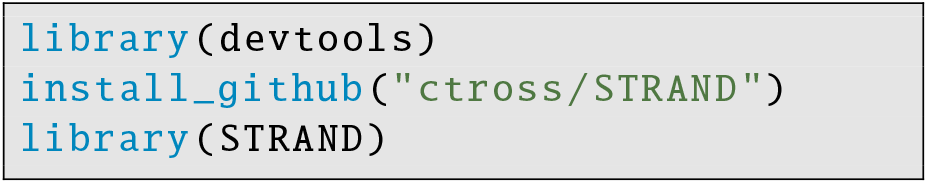

All of the tutorial code elaborated on below can also be found online at: https://github.com/ctross/STRAND, where the package will be maintained.

### Building data objects

The first step in building any `STRAND` model is to organise the data. Social network data are normally complex, with some variables being reported at the level of the individual and others being reported at the level of the dyad. The `make strand data` function serves to organise all of these data into a unified format that can be read by later functions. After data are compiled, they can then be analysed with simple, `lm`-style function calls, as we discuss below.

We will illustrate how `STRAND` data objects are built, using human friendship network data from Dalla Ragione et al. (2022). First, outcome data and dyad-level predictors (both structured as adjacency matrices) are stored as labeled lists:

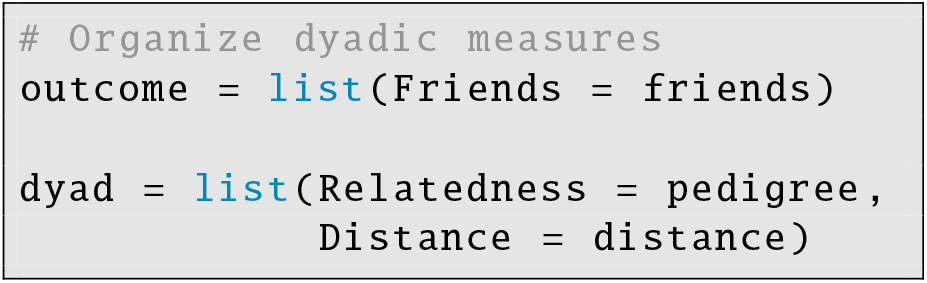

The outcome data must be either binary data or integers. The dyadic covariate data can include numeric variables, indicator variables, or even categorical variables. These dyadic covariate data can be used to estimate associations between dyad-level characteristics—such as genetic relatedness or physical proximity—and the likelihood of a tie in the outcome network.

Next, the individual-level covariates are stored in a dataframe:

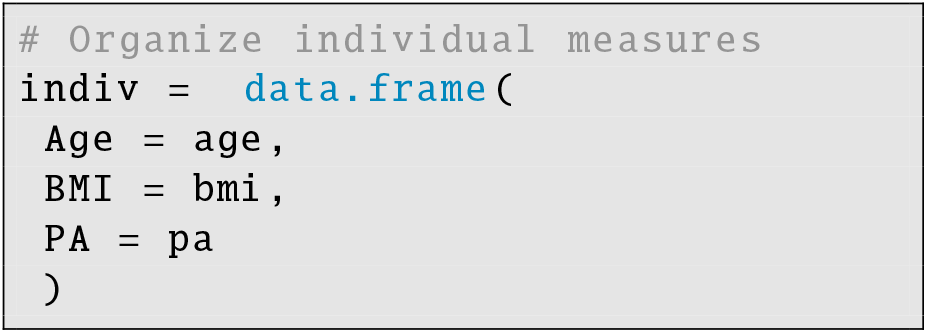

The individual-level covariate data can also include numeric variables, indicator variables, or categorical variables. Individual-level covariate data can be used to estimate associations between individual-level characteristics and the likelihood of either sending or receiving a tie.

Finally, individual-level covariates that govern group/block structure are stored in a separate data-frame:

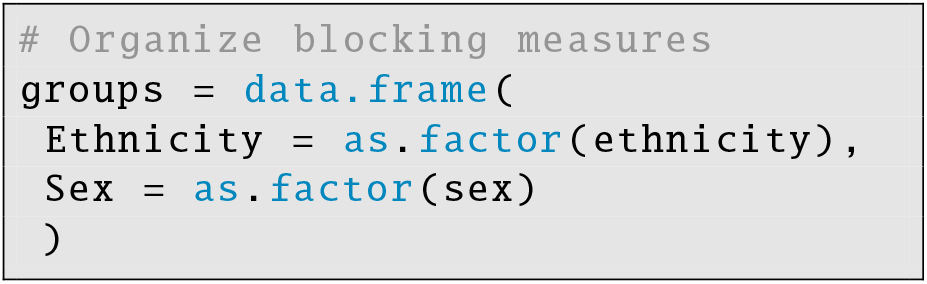

Although these block-structuring variables are also individual-level data, they are treated differently than other variables by `STRAND`; these variables must be factors, and are used to create random intercept offsets unique to the interaction of focal/sender and alter/receiver block IDs.

Once all covariate data are organised as above, they can be compiled into a single `STRAND` object:

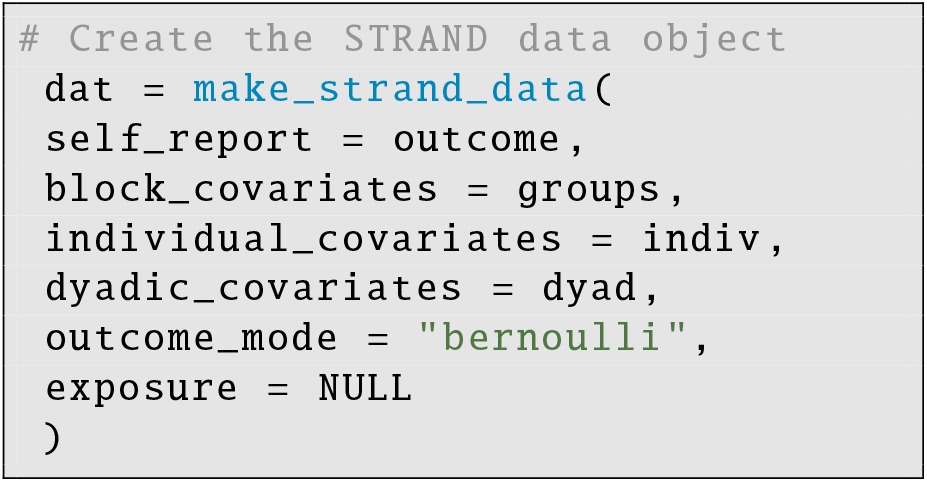

At this point, the user must define which outcome model to use. The `STRAND` package supports three outcome modes for each model type: “bernoulli” for binary tie data (e.g., for human self-report/name-generator data, or similar binary tie data from non-human animals, like whether or not there was any observed conflict between each dyad on a given day), “poisson” for raw count data (e.g., the number of times GPS trackers were within 5 meters of each other over a fixed 1-week period), or finally “binomial” for proportion data (e.g., if the outcome variable is a matrix containing a count of grooming events between each dyad, and the exposure variable is matrix containing a count of the number of scans in which grooming events between each dyad could have been observed). If the outcome mode is set to “binomial”, then the `exposure` variable (a labeled list containing a matrix of sample size values) must be provided.

### Example model with Binary data

Classic social network data, especially as collected using self-report survey designs in human research, is frequently represented as a matrix of binary ties (i.e., zeros indicating the absence of ties, and ones indicating the presence of ties). For our first example, we will model binary human friendship network data (see Figure 1) published in Dalla Ragione et al. (2022). These data include individual-level measures (e.g., sex, ethnicity, age, BMI, and physical attractiveness) and dyad-level measures (e.g., relatedness and interhousehold distance) that are thought to be associated with the structure of friendships.

**Figure 1.**
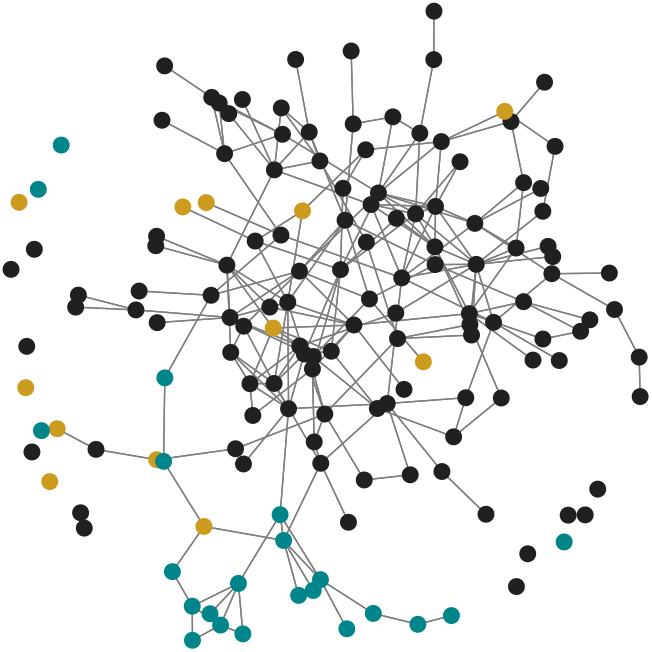
Friendship network data from a rural Colombian town published in Dalla Ragione et al. (2022). Nodes are coloured according to ethnic group, with blue nodes representing indigenous Emberá individuals, goldenrod nodes representing Mestizo individuals, and dark-grey nodes representing Afrocolombian individuals. Group structure is modelled by including ethnicity as an observed block variable. Variance in node degree is estimated using random effects on the probability of sending and receiving friendship nominations.

To model the data, we use a hybrid of the stochastic block-model and social relations model (see Supporting Information for full mathematical details). The `STRAND` syntax is based on standard `lm` syntax from base R. To model the data, we write out equations for block effects, focal/sender effects, alter/receiver effects, and dyadic effects:

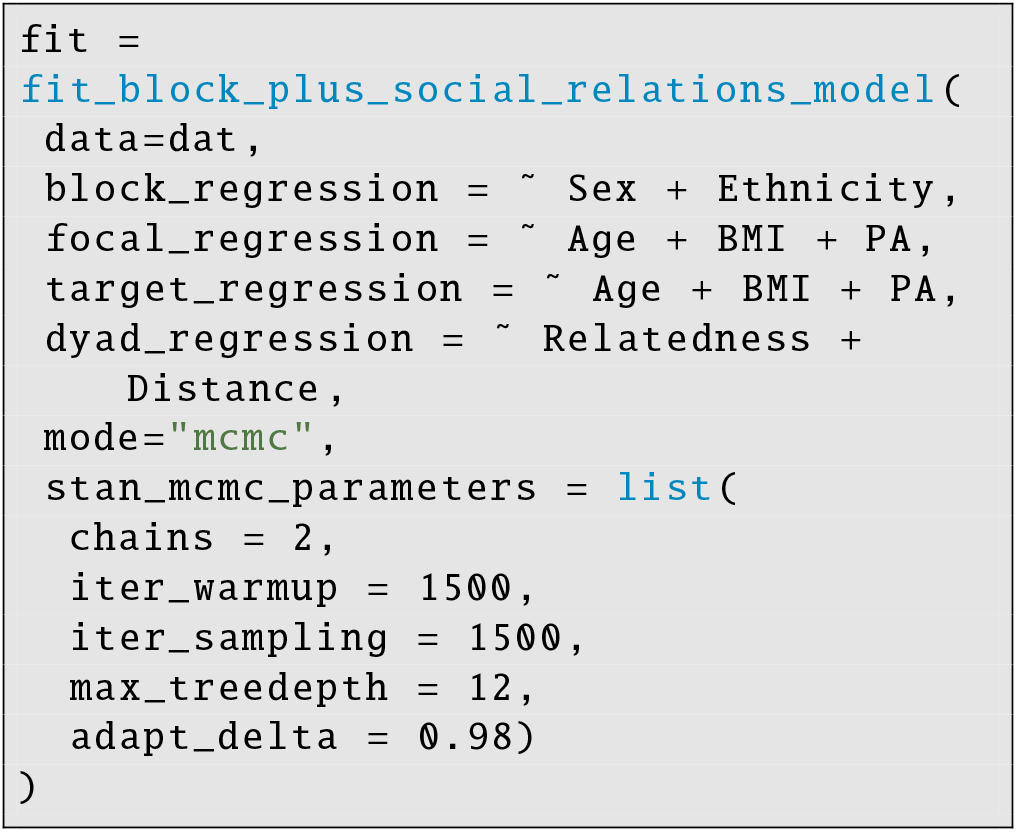

In this model, we estimate block-level effects for sex and ethnicity using the argument: `block_regression =` ∼`Sex + Ethnicity`. These effects are indicative of how the likelihood of friendship nomination varies as a function of block categories—i.e., we can measure if male-to-male ties are more or less likely than male-to-female ties, female-to-male ties, or female-to-female ties (and likewise for ties within and between ethnic groups).

Next, the focal regression model, `focal_regression =` ∼ `Age + BMI + PA`, explores how the age, body mass index, and physical attractiveness rating for a given individual is related to that individual’s propensity to nominate others as friends (i.e., it measures the effects of individual-level covariates on *out-degree*). Similarly, the target regression model, `target_regression =` ∼ `Age + BMI + PA`, explores how the age, body mass index, and physical attractiveness rating for a given individual is related to that individual’s propensity to be nominated by others as a friend (i.e., it measures the effects of individual-level covariates on *in-degree*). Finally, the dyad regression model, `dyad_regression =` ∼ `Relatedness + Distance`, explores how the likelihood of friendship ties is associated with the genetic relatedness of each dyad, as well as with the spatial distance between the homes of each dyad.

When the model has finished running, the MCMC samples can be processed and summarised using a convenience function:

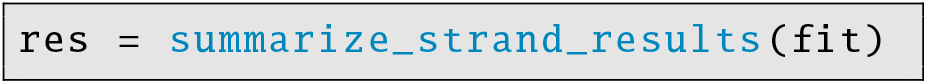

The results can then be visualised as a table (e.g., see Table 1) or as a forest plot (e.g., see Figure 2).

**Table 1.**
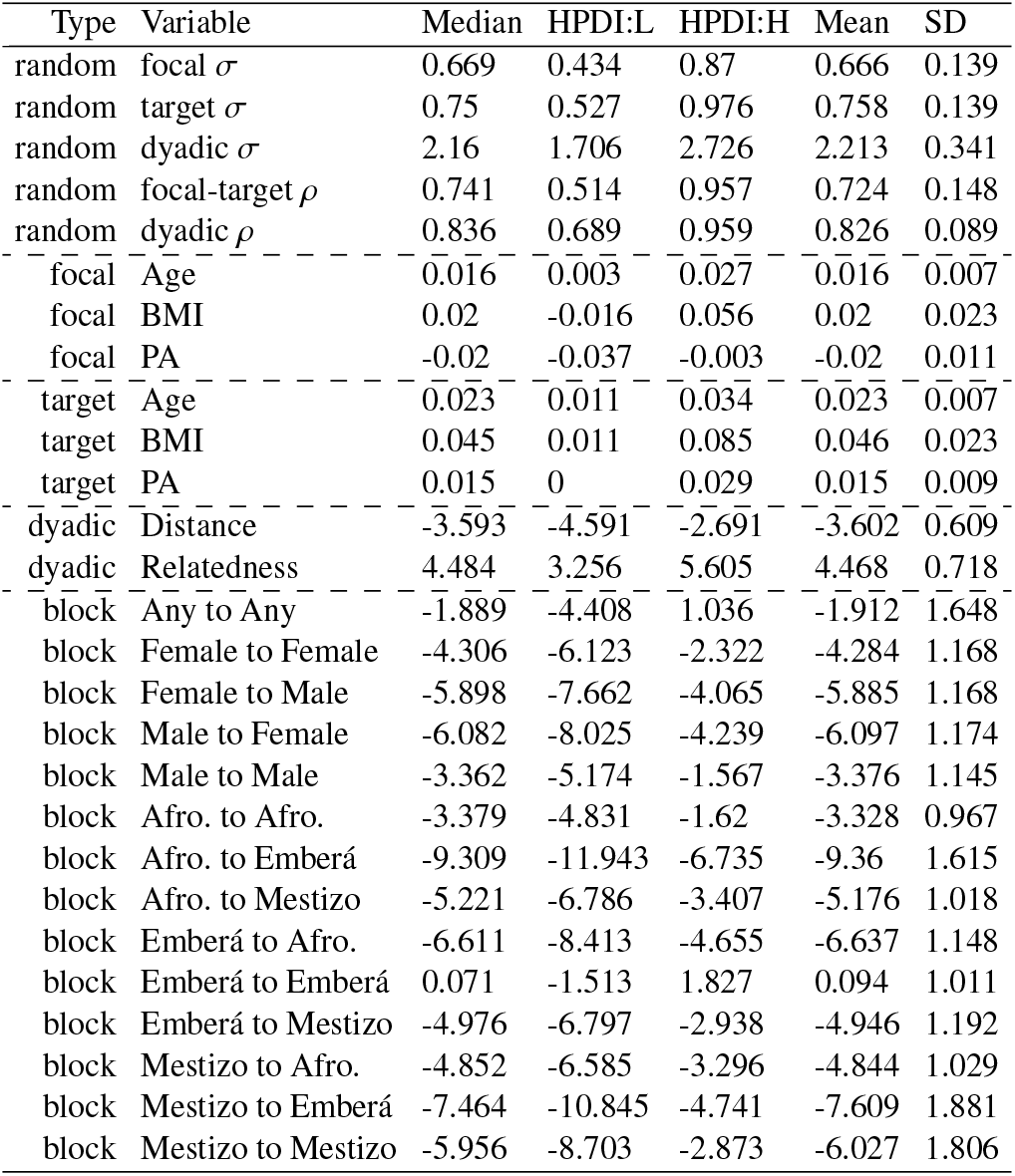
Parameter estimates for the effects of various covariates on the structure of friendships in a rural Colombian town. *Focal, target*, and *dyadic* effects are interpreted as slopes. *Block* effects are interpreted as intercept offsets. *Random* effects include terms that control the variance of random effects, *σ*, and terms that control the correlation of random effects, *ρ*.

**Figure 2.**
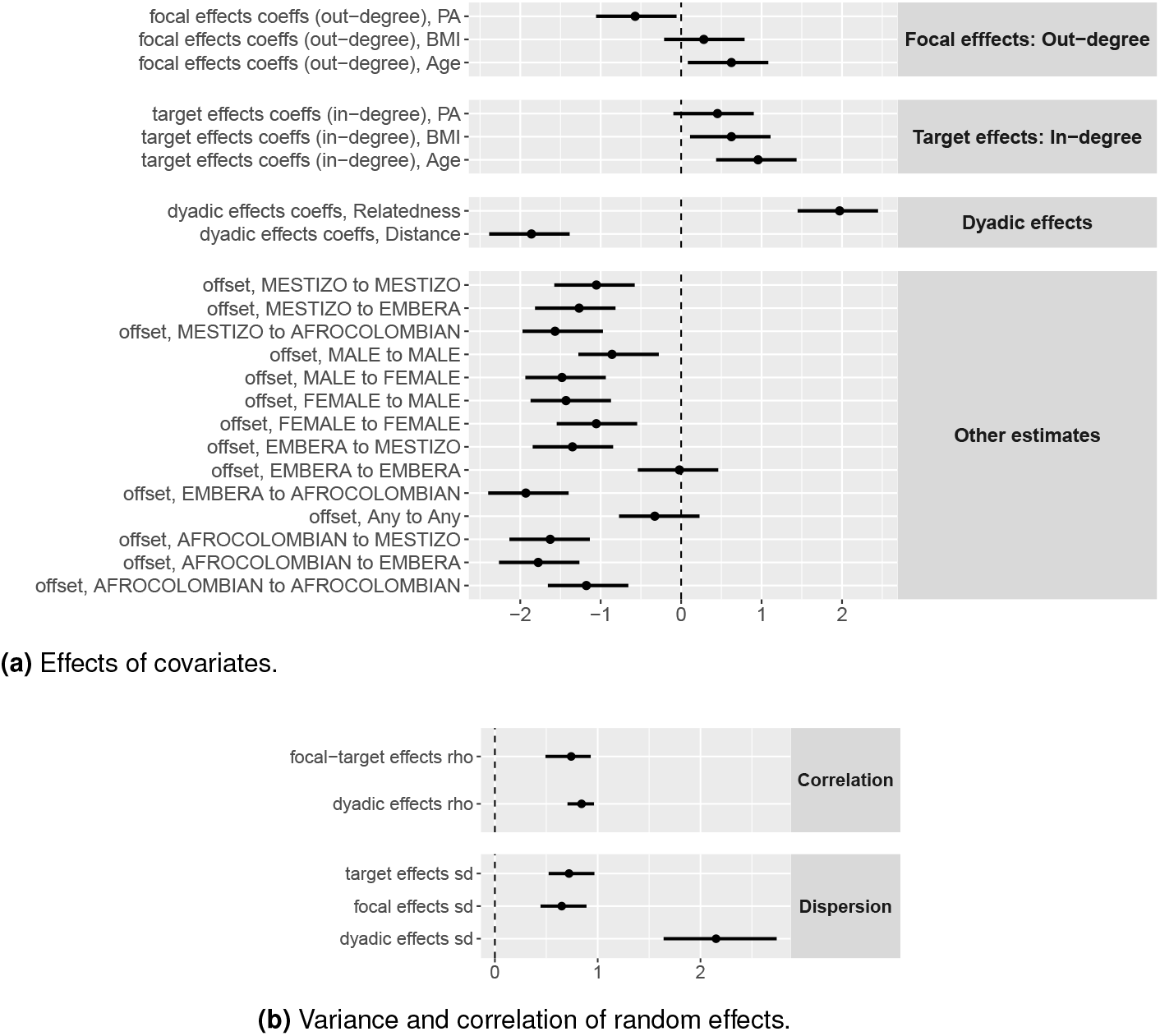
Parameter estimates for the effects of various covariates on the structure of friendship in a rural Colombian town. Points represent the posterior median, and bars represent 90% highest posterior density intervals.

As in Dalla Ragione et al. (2022), we find that relatedness is positively associated with the probability of observing friendship ties, while inter-household distance is negatively associated the probability of friendship ties. Older individuals have a higher probability of making friendship nominations, while individuals who were rated by the community as more physically attractive are less likely to nominate others as friends. On the other hand, individuals rated as more physically attractive were more likely to be nominated by others as friends. Age and BMI are also associated with the likelihood on being nominated as a friend. Next, when considering block variables, such as ethnicity and sex, we find evidence that individuals preferentially assort with others of their same group. For example, the intercept offset for Afrocolombian-to-Afrocolombian ties is -3.37 (90%HPDI; -4.83, -1.62), is reliably greater than the intercept offset for Afrocolombian-to-Emberá ties, -9.30 (90%HPDI; -11.94, -6.73). Finally, we find evidence of both dyadic reciprocity (dyadic *ρ*) and generalised reciprocity (focal-target *ρ*) in friendship nominations (see Table 1).

### Example model with Poisson data

Throughout the non-human literature, network data are often recorded using numerical measurements (e.g., the number of times two animals are observed fighting, or the number of minutes two animals spend grooming one another). As an example of such numerical measures, we draw on data investigating blood-sharing among vampire bats (Carter and Wilkinson 2013). These data include individual-level predictors (i.e., sex) and dyad-level predictors (i.e., genetic relatedness, and whether each dyad had the opportunity to be observed sharing blood).

As before, we start by organising the data:

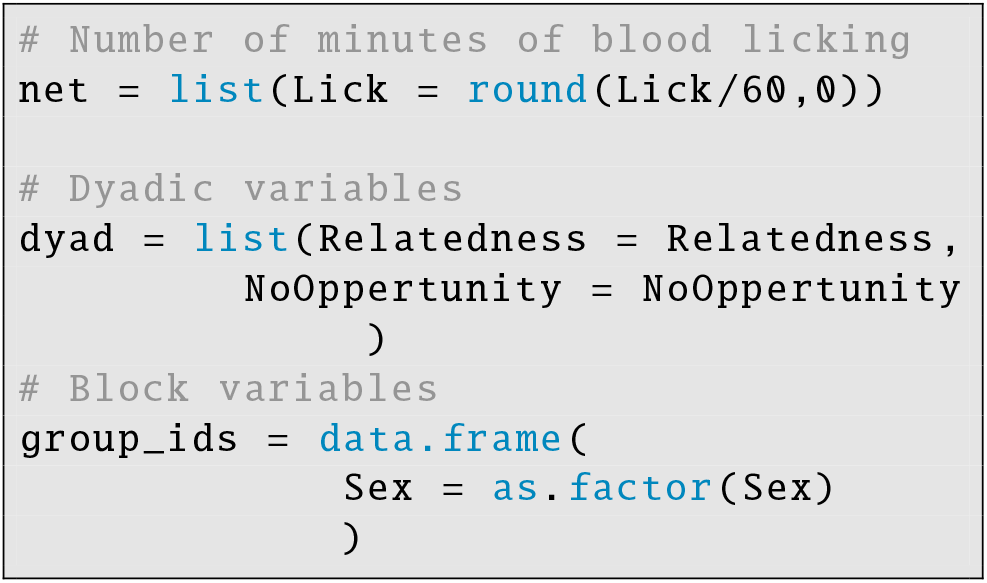

Then, the data can be compiled into a `STRAND` data object. This time, we must include the argument: `outcome_mode = “poisson”`, so that `STRAND` treats the outcome data as integers. Also, since there are no individual-level covariates other than sex, which is used as a blocking variable, we can set: `individual_covariates = NULL`.

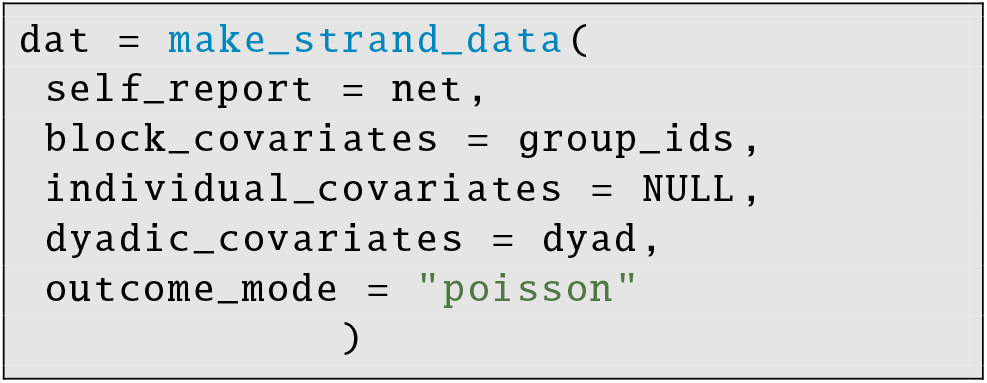

Then, a model can be fit to the data:

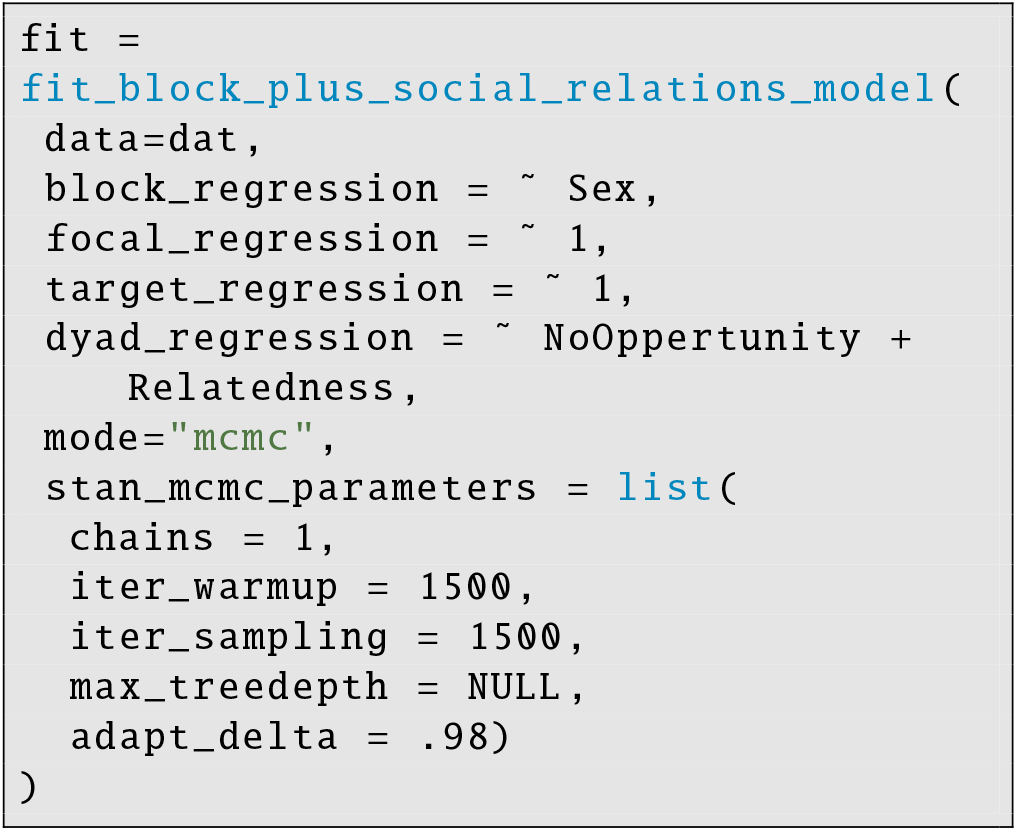

Here, we set the focal and target regression models to be intercept only (as we have no individual-level covariates) using the `lm`-style syntax: `focal_regression =` ∼ `1` and `target_regression =` ∼ `1`.

Finally, the results can be summarised and plotted:

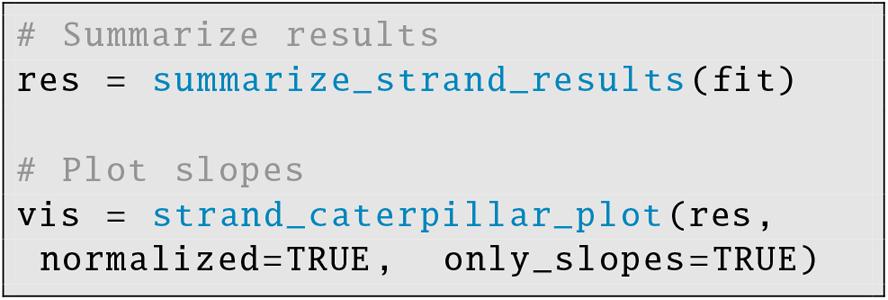

Table 2 and Figure 4 present the effects of relatedness and sex on the rate of blood-sharing transfers. We recover the primary results of Carter and Wilkinson (2013), finding that genetic relatedness is a reliable predictor of blood-sharing, and that transfers are reliably more likely between female dyads than between male or mixed-sex dyads.

**Table 2.** Predictors of vampire bat blood-sharing relationships.

| Type | Variable | Median | HPDI:L | HPDI:H | Mean | SD |
| --- | --- | --- | --- | --- | --- | --- |
| random | focal $\sigma$ | 0.855 | 0.322 | 1.477 | 0.872 | 0.348 |
| random | target $\sigma$ | 1.3 | 0.667 | 2.05 | 1.332 | 0.437 |
| random | dyadic $\sigma$ | 2.475 | 2.09 | 2.977 | 2.496 | 0.274 |
| random | focal-target $\rho$ | 0.23 | -0.32 | 0.72 | 0.196 | 0.322 |
| random | dyadic $\rho$ | 0.62 | 0.449 | 0.804 | 0.61 | 0.113 |
| dyadic | No Opportunity | -0.971 | -2.016 | 0.207 | -0.976 | 0.685 |
| dyadic | Relatedness | 1.299 | 0.121 | 2.426 | 1.285 | 0.726 |
| block | Any to Any | 0.716 | -1.213 | 2.713 | 0.694 | 1.216 |
| block | Female to Female | -0.275 | -2.164 | 1.819 | -0.29 | 1.248 |
| block | Female to Male | -4.203 | -6.08 | -2.228 | -4.209 | 1.226 |
| block | Male to Female | -3.607 | -5.427 | -1.513 | -3.563 | 1.226 |
| block | Male to Male | -6.439 | -8.481 | -4.151 | -6.421 | 1.331 |

**Figure 3.**
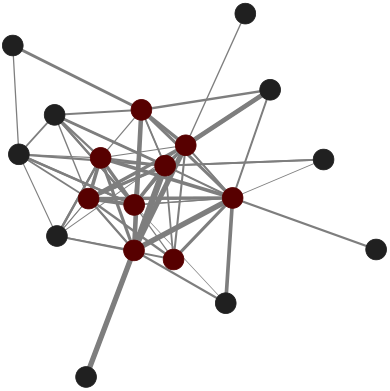
Blood-sharing network data from vampire bats published by Carter and Wilkinson (2013). Red nodes represent females and dark-grey nodes represent males. Group structure is modelled by including sex as a block variable. Variance in node degree is estimated using random effects.

**Figure 4.**
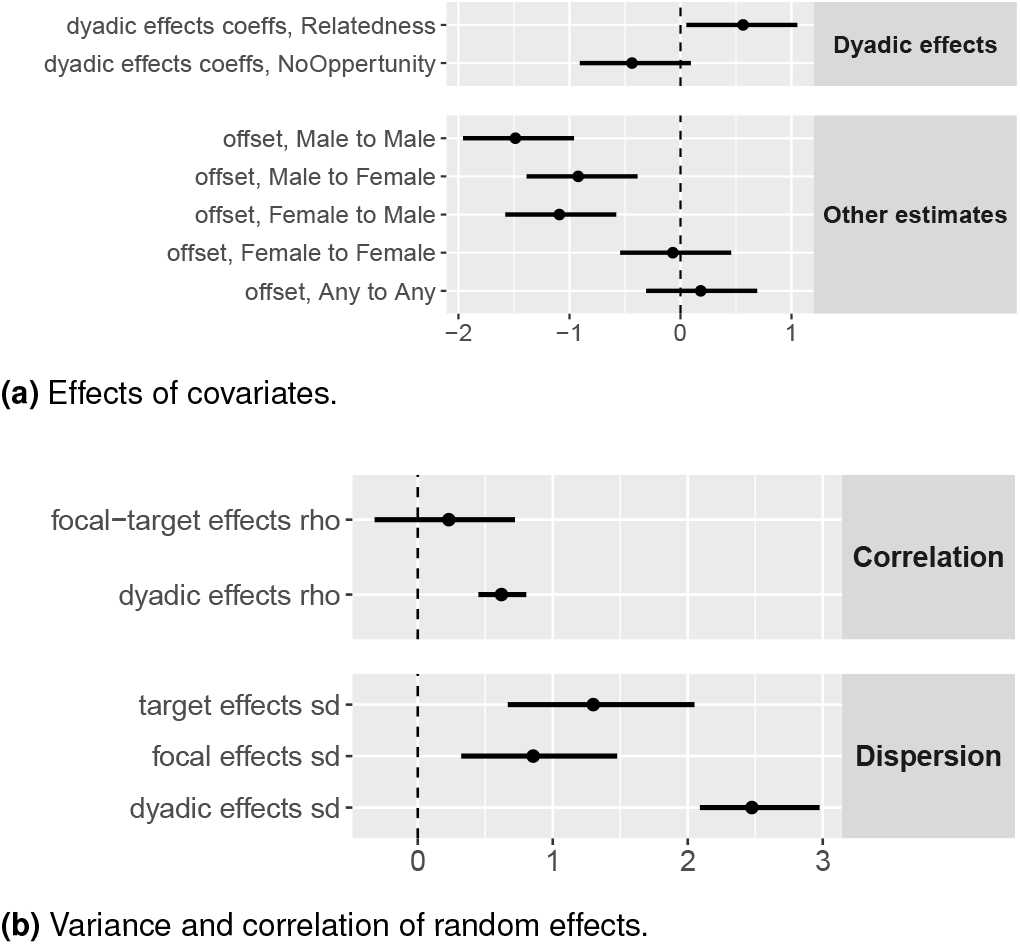
Blood sharing among of vampire bats. Points represent the posterior median, and bars represent 90% highest posterior density intervals. Vampire bats are more likely to share blood with relatives than less-related conspecifics. Transfers are also more likely to flow from females to other females than from females to males, as indicated by the credible regions for these offsets not overlapping. Blood sharing is also reciprocal, as indicated by the strong correlation in dyadic random effects.

### Example model with Binomial data

Another type of data frequently encountered in studies of animal sociality represents ties strength as a weighted social association matrix (e.g., Brask et al. 2019). These measures are typically created by applying a simple ratio association index (SRI; Cairns and Schwager 1987; Farine and Whitehead 2015), where, for example, the edge weight of each dyad is calculated as the ratio of the number of scans/observations in which the dyad is observed together divided by the number of scans/observations in which at least one of them was observed.

While this approach of weighting counts by an exposure variable is significantly better than ignoring variation in risk of observation (Farine and Whitehead 2015), construction of a simple ratio divides out sample size information, leading dyadic observations based on little data to carry disproportionate weight in downstream analyses (see McElreath 2020; Hart et al. 2021b, for a review of this issue). Moreover, zeros arising from censoring (i.e., due to members of a dyad being unavailable; “denominator zeros”) are often confounded with true zeros (i.e., members of a dyad being present but not interacting; “numerator zeros”). A better approach involves modelling the actual *count* of the number of scans/observations in which each dyad is observed together using a Binomial model, in which the sample size parameter is—for example—the number of scans/observations in which at least one member of the dyad was observed.

To demonstrate how to fit such a model in `STRAND`, we draw on grooming data from captive Guinea baboons published by Gelardi et al. (2020). We investigate three questions here: 1) do individuals who groom others more also receive more grooming in general (i.e., generalised reciprocity)?, 2) accounting for individual-level differences in the probability of grooming, does the probability of individual *i* grooming individual *j* increase with the probability that individual *j* grooms individual *i* (i.e., dyadic reciprocity)?, and 3) is the probability of individual *i* grooming individual *j* associated with whether individual *j* “presents” to individual *i*? (Presenting is defined as: “*approaching another individual gently with or without lipsmacks and grunts and presenting the rear*”; Gelardi et al. 2020).

As before, we start by organising the data:

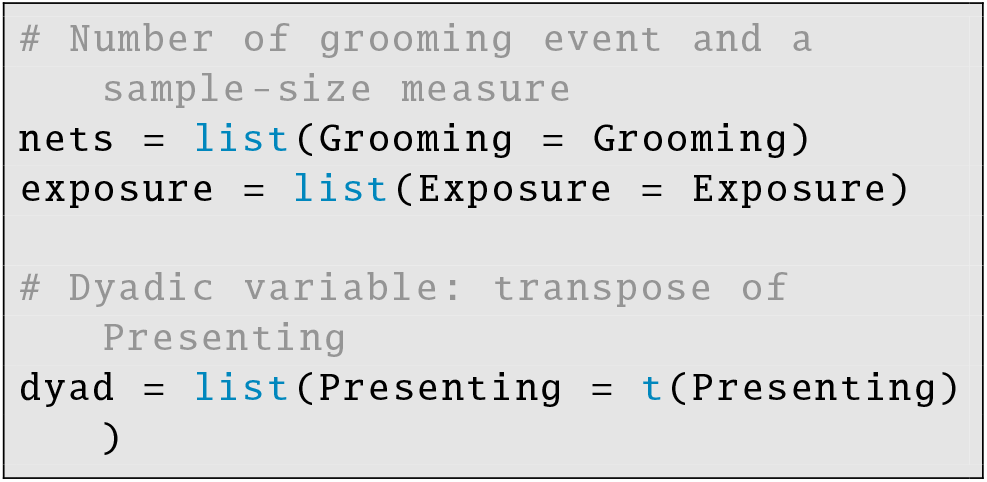

And then we compile the data into a `STRAND` object:

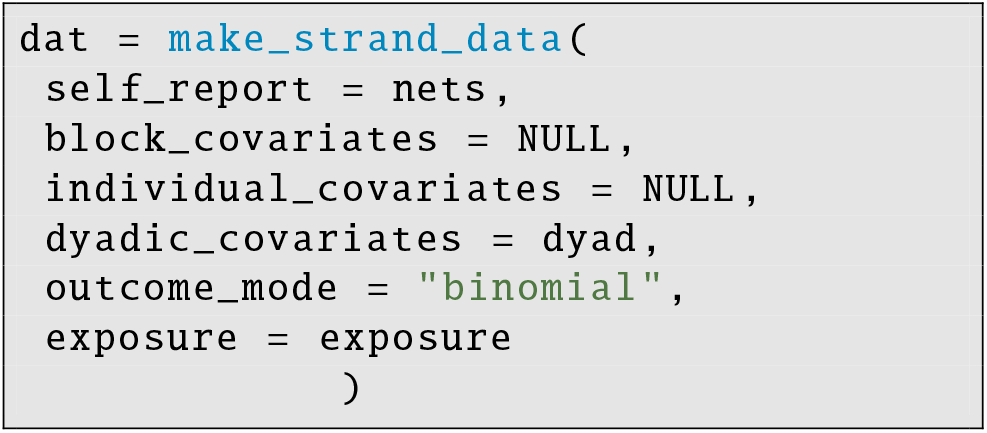

This time, the argument: `outcome_mode = “binomial”` is included. In order for the Binomial model to run, we also need to store the sample size data, using: `exposure = exposure`.

This time, because we have no blocking variables, we use the basic social relations model with no block-level effects. We run this model using the function:

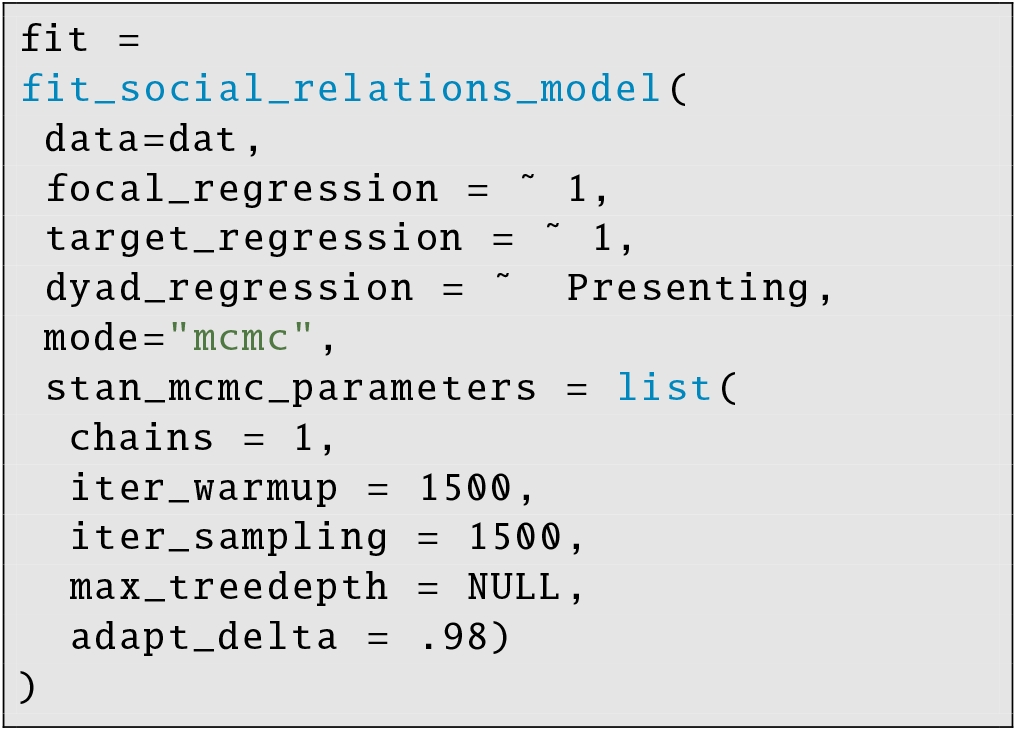

Finally, the results can be summarised and plotted:

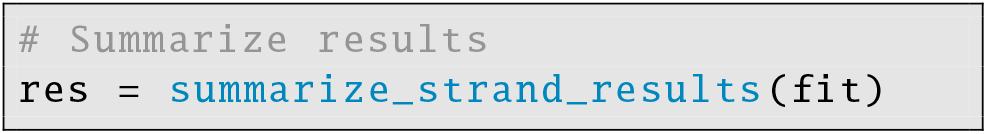

The results are presented in Table 3 and Figure 6. We note that the correlation, *ρ*, of focal and target effects is negative. This indicates that baboons who frequently groom others are actually less likely to be groomed themselves by others. Such unidirectional behavioural propensities are consistent with dominance hierarchies (e.g., Gullstrand et al. 2021) in which unbalanced benefits are tolerated. However, after accounting for this individual-level variation in grooming propensity, there is evidence of dyadic reciprocation, as indicated by the strong correlation, *ρ*, in dyadic random effects. Finally, in this sample of Guinea baboons, individuals appear more likely to groom conspecifics who regularly “present” to them.

**Table 3.** Results of captive Guinea baboon grooming.

| Type | Variable | Median | HPDI:L | HPDI:H | Mean | SD |
| --- | --- | --- | --- | --- | --- | --- |
| random | focal $\sigma$ | 0.865 | 0.529 | 1.242 | 0.896 | 0.234 |
| random | target $\sigma$ | 0.541 | 0.266 | 0.882 | 0.559 | 0.195 |
| random | dyadic $\sigma$ | 1.455 | 1.233 | 1.682 | 1.46 | 0.138 |
| random | focal-target $\rho$ | -0.483 | -0.878 | -0.068 | -0.444 | 0.266 |
| random | dyadic $\rho$ | 0.603 | 0.369 | 0.779 | 0.59 | 0.128 |
| dyadic | Any to Any | -5.645 | -6.161 | -5.225 | -5.66 | 0.294 |
| dyadic | Presenting | 0.339 | 0.159 | 0.491 | 0.341 | 0.103 |

**Figure 5.**
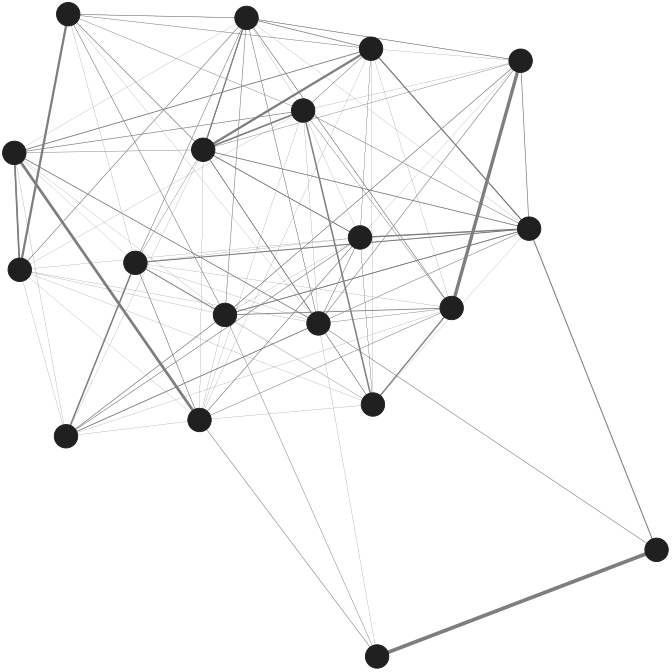
Grooming data in captive Guinea baboons published by Gelardi et al. (2020). Group structure is not modelled as no individual-level data were provided. Variance in node degree is estimated using random effects on the probability of providing and receiving grooming.

**Figure 6.**
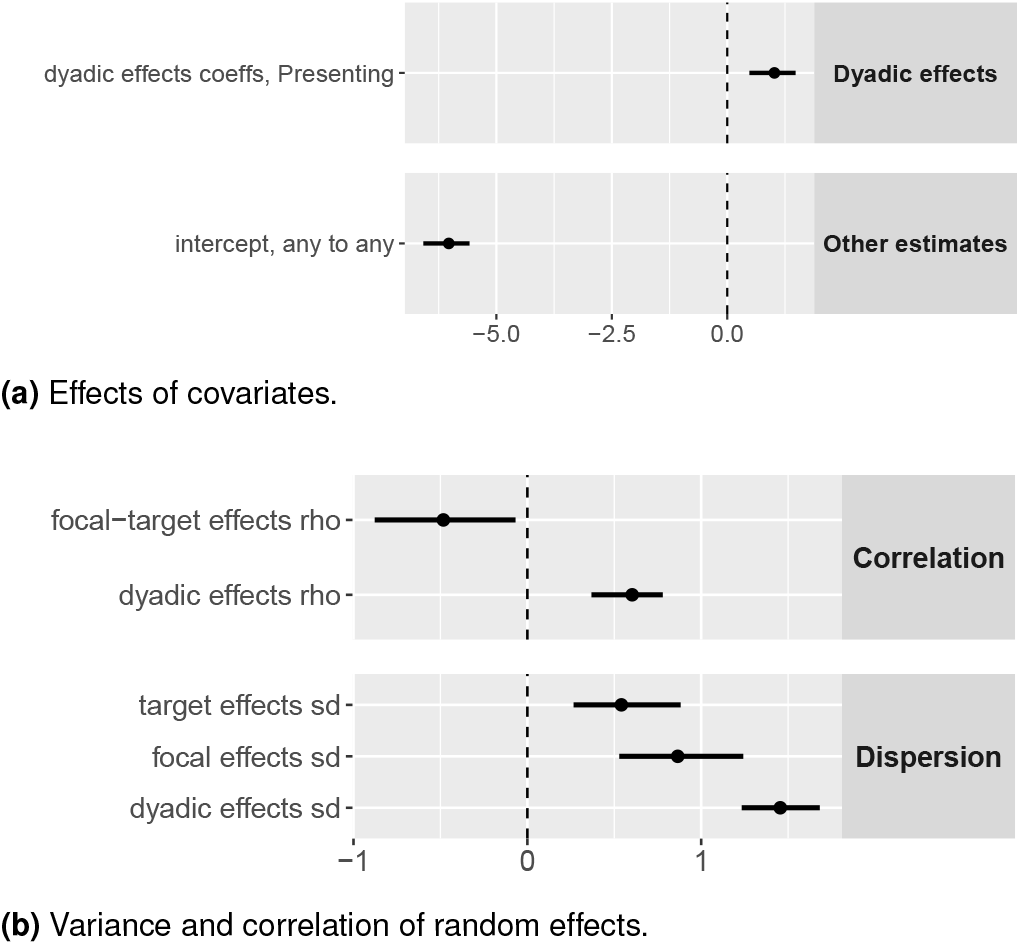
Grooming data in captive Guinea baboons. Points represent the posterior median, and bars represent 90% highest posterior density intervals. Guinea baboons appear more likely to groom conspecifics who regularly “present” to them. Interestingly, the correlation, *ρ*, of focal and target effects is negative. This indicates that baboons who frequently groom others are less likely to be groomed themselves by others. However, after accounting for this individual-level variation in grooming propensity, there is evidence of dyadic reciprocation, as indicated by the strong correlation, *ρ* in dyadic random effects.

### Vaildating the models with simulated data

For interested readers, we include a detailed mathematical description of our statistical models in the Supporting Information. There, we walk readers through model specification, parameter interpretation, and model validation procedures. We test each statistical model, for each outcome mode (Bernoulli, Poisson, and Binomial) using a suite of unit-tests (see also Redhead et al. 2021). Specifically, we first generate network data using forward simulations from a stochastic blockmodel, a social relations model, or the combined model, which includes both stochastic blockmodel and social relations model parameters. We then use the corresponding inferential statistical models to analyse the simulated data sets and ensure that we can recover the generative parameter values. In each simulation experiment, we generally vary only a single generative parameter (e.g., the dyadic reciprocity coefficient) across a broad parameter space that contains realistic values, while fixing all other parameters in the model to empirically plausible values. See Figure 7 for an example, and the Supporting Information for the full suite of unit-tests, which all indicate that our models accurately recover generative parameters.

**Figure 7.**
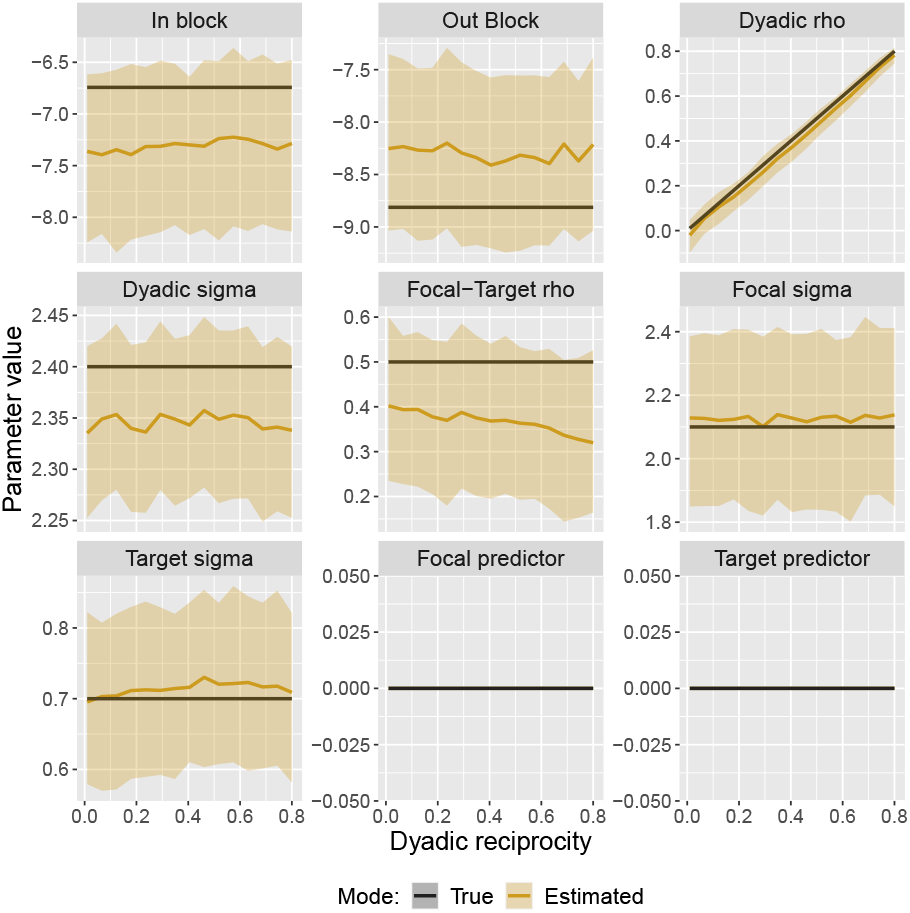
Example of model validation results. In this case, we vary the dyadic reciprocity parameter of a generative network model and simulate test data-sets for a range of values, *ρ*_*δ*_ ∈ (0.01, 0.8). We then use STRAND to estimate the model. Black lines represent the generative parameter values. Yellow regions represent estimated posterior distributions of the same parameters. We find that our model accurately recovers all generative parameters. We repeat this process for all combinations of network models and outcome modes, varying all key model parameters for each test case. See supporting information for details.

## Conclusions

The tools included in `STRAND` provide easy-to-use and efficient methods for generative modelling of human and animal social networks. Here, we have outlined the functionality of `STRAND`, defined the suite of models that are included, and provided detailed unit-tests to show that the software performs correctly. Using openly available example data-sets, we have provided tutorials for endusers interested in running network analysis models in R using `STRAND`. We hope that this software will help end-users with limited programming experience easily deploy otherwise complex statistical models, and thus effortlessly investigate fundamental research questions in their fields of interest. End-users can find a complete index of—and full documentations for—all functions included in `STRAND` by visiting: https://github.com/ctross/STRAND. Additional R code examples are provided there as well.

## Acknowledgements

C.T.R., R.M.E., and D.R. were supported by the Department of Human Behavior, Ecology, and Culture at the Max Planck Institute for Evolutionary Anthropology. Leipzig, Germany. D.R. was supported by ESRC Grant number ES/V006495/1. Authors declare no conflict of interest.

