## Supplementary Materials for "Modelling human and non-human animal network data in R using STRAND"

*Submitted to: Submitted to Methods in Ecology & Evolution*
May 6, 2022

### Supplementary Materials to: *Modelling human and non-human animal network data in R using STRAND*

Cody T. Ross[1], Richard McElreath[1], and Daniel Redhead[1*]

[1] Department of Human Behaviour, Ecology and Culture
Max Planck Institute for Evolutionary Anthropology, Deutscher Platz 6, 04103 Leipzig, Germany.

#### 1. Model definition

In Redhead et al. (2021), we first introduced the `STRAND` R package for analysing human network data as collected via the 'name-generator' method. The package was initially focused only on modelling double-sampled, binary, self-report data. However, across the social and biological sciences, researchers often have single-sampled data—which may be represented as binary indicators, counts, or proportions. As such, in order to make the `STRAND` package more useful to a wider-variety of researchers, we have added support for single-sampled networks. We have also revised the supported models to allow multiple block memberships (i.e., to allow for block-level effects of multiple covariates, such as sex and social group, simultaneously). We have further added support for non-Bernoulli outcomes: `STRAND` supports count data through Poisson models, and proportion data through Binomial models. In the following subsections, we review the full statistical specification for these three families of models. After this, we provide the results of a suite of unit-tests, in which we fit each model to simulated data. Through this, we have ensured that each statistical model is capable of recovering the parameters of a generative model used to simulate testing data.

##### *1.1. The standard Bernoulli model*

`STRAND` was originally designed to model Bernoulli outcomes produced under stochastic blockmodels, social relations models, and a hybrid of these two models. We review the general structure of these models below, and show how they are adapted to Poisson and Binomial outcomes in the following sections.

###### 1.1.1. Block or community structure

Across many real-world networks, individuals within a given network may form tight-knit subgroups (blocks), and there may be a higher probability of ties forming within these blocks. Such subgroups may be formed on the basis of easily observable attributes—such as rank or sex. The extent of this partitioning of individuals into sub-groups may be estimated using a stochastic blockmodel design (Pearl and Schulman 1983; Wasserman et al. 1994). Here, we assume that block membership for each individual is known data, and that individuals belong to a single block in each of the $V$ observable group/block variables.

Diverging slightly from the notation used in Redhead et al. (2021), we let the adjacency matrix, $Y$, denote an observed network of ties. The elements $Y_{[i,j]} \in \{0, 1\}$ reflect the presence or absence of directed ties (e.g., resource transfers, or aggression events) from individual $i$ to individual $j$. For each grouping variable $v \in \{1, ..., V\}$, the community of $N$ individuals is assumed to be divisible into $K_{[v]}$ blocks, or sub-communities. We further assume that each individual belongs to only one block for each variable, $v$, that the block of type $k$ for variable $v$ has sample size $N_{[k,v]}$, and that the block of individual $i$ in variable $v$ is returned by the function $b(i, v)$. We let the elements of the square matrix, $B_{[v]}$, denote the log-odds offsets of tie likeliness between individuals in various blocks. To generate an intercept term, we let $B_{[0]}$ be defined as a one-by-one matrix, and we let $b(i, v = 0) = 1$ for all $i$.

Suppose that we have a community structure where there are two grouping variables, $V = 2$. Next, suppose that for the first variable we have three different kin-group labels (e.g., we can classify individuals as belonging to one of three distinct kin groups); so $K_{[1]} = 3$. For the second variable, suppose that we have two rank groups (e.g., we can classify individuals as being high-rank or low-rank); so $K_{[2]} = 2$.

The first $B_{[v]}$ matrix could then be written as:

$$B_{[1]} = \begin{bmatrix} \beta_{1\to1} & \beta_{1\to2} & \beta_{1\to3} \\ \beta_{2\to1} & \beta_{2\to2} & \beta_{2\to3} \\ \beta_{3\to1} & \beta_{3\to2} & \beta_{3\to3} \end{bmatrix} \tag{1}$$

and the second as:

$$B_{[2]} = \begin{bmatrix} \beta_{1\to1} & \beta_{1\to2} \\ \beta_{2\to1} & \beta_{2\to2} \end{bmatrix} \tag{2}$$

The parameters on the diagonal within each matrix control the probabilities of within-group ties, and the parameters on the off-diagonals control the probabilities of between-group ties. For instance, if ties tend to flow in one direction—e.g., from individuals in group 1 to individuals in group 2—then the parameters in one direction may be larger than in the opposite direction—e.g., $\beta_{1\to2} > \beta_{2\to1}$. Note that, although we abuse notation a bit for visual clarity, the parameter labeled $\beta_{1\to2}$ in $B_{[1]}$ is a distinct parameter from $\beta_{1\to2}$ in $B_{[2]}$.

Assuming only this block-induced sub-structure, the generative model for $Y_{[i,j]}$ may be written as:

$$Y_{[i,j]} \sim \text{Bernoulli}\left(\text{Logistic}\left(\sum_{v=0}^{V} B_{[v,b(i,v),b(j,v)]}\right)\right) \tag{3}$$

where the probability of a tie from individual $i$ in block $b(i, v)$ to individual $j$ in block $b(j, v)$ for variable $v$ is controlled by the corresponding entry in the array of block parameters, $B[v, b(i, v), b(j, v)]$.

In typical cases, the diagonal elements of $B_{[v]}$, which control the frequency of ties within a block, will have higher prior weight than the off-diagonal elements, though other topologies are possible (see Batagelj 1997). Alongside this, priors should also depend on sample size, $N$, so that the prior network density has empirically plausible scaling over a wide range of $N$ values. The priors specified below accomplish both of these goals, and were selected on the basis of robusticity checks (see details in Redhead et al. 2021):

$$\beta_{k\to k} \sim \text{Normal}\left(\text{Logit}\left(\frac{0.1}{\sqrt{N_{[k,v]}}}\right), 1.5\right) \tag{4}$$

$$\beta_{k\to\tilde{k}} \sim \text{Normal}\left(\text{Logit}\left(\frac{0.01}{0.5\sqrt{N_{[k,v]}}+0.5\sqrt{N_{[\tilde{k},v]}}}\right), 1.5\right) \tag{5}$$

Here, $k \to k$ indicates a diagonal element and $k \to \tilde{k}$ indicates an off-diagonal element. The scalar of 0.1 in Eq. 4 places higher prior density on the diagonal of $B$ (which controls the probability of within-block ties), than the off-diagonal of $B$ (where the scalar of 0.01 from Eq. 5 generates reduced prior between-block tie probability). The scalars of $\sqrt{N_k}$ ensure that prior tie probability scales with sample size at roughly the same rate that we see in empirical data sets (Ready and Power 2021). The standard deviation of 1.5 in both equations causes the overall prior to be rather weak, and thus allows the data to dominate the posterior. These priors, and all other default priors discussed below, however, can be modified by the user according to their needs by passing in a labeled list of priors when calling the model function.

##### 1.1.2. The social relations model

Alongside the gross topologies produced by the stochastic block model, many social networks are characterised by individual-level variation in the propensity to form ties. For example, male vampire bats (in the second example in the main text) were unlikely to share blood (Carter and Wilkinson 2013). Many social networks exhibit a right-skewed distribution of ties (i.e., a 'degree distribution' with a fat upper tail)—such that many individuals have relatively few ties, while a small number of individuals have a large number of ties (Barabási and Albert 1999; Broido and Clauset 2019; Newman 2001). Given this, we incorporate correlated individual-level random effects (i.e., 'sender' and 'receiver' effects) that control the propensity of nodes to send outgoing, and receive incoming, ties. The correlation between these random effects can further be used to measured "generalised reciprocity", as seen in the third example of the main text on baboon grooming behaviour (Gelardi et al. 2020).

Specific pairs of individuals (i.e., dyads) within any given network may also be more or less likely than chance to share mutual ties. While community structure (outlined above) provides a mechanistic explanation for some degree of reciprocity, models accounting for community structure alone typically fail to produce a realistic amount of reciprocity (see, Safdari et al. 2021). As such, we include dyadic random effects to permit higher levels of dyadic reciprocity (Snijders and Kenny 1999), above and beyond that which is explained by community structure alone. In all three of our example models, in both humans and non-human mammals, we found evidence of such "dyadic reciprocity".

By integrating the SRM and the SBM approaches, Eq. 3 may be replaced by Eq. 6:

$$Y_{[i,j]} \sim \text{Bernoulli}\Big(\text{Logistic}(\phi_{[i,j]})\Big) \tag{6}$$

where:

$$\phi_{[i,j]} = \left( \sum_{v=0}^{V} B_{[v,b(i,v),b(j,v)]} \right) + \lambda_{[i]} + \pi_{[j]} + \delta_{[i,j]} + \dots \tag{7}$$

Here, $B$ is the list of SBM matrices (which control the tendency for ties to form within and between blocks), $\lambda$ is a vector of individual-specific sender/nominator effects governing out-degree, $\pi$ is a vector of individual-specific receiver/target effects governing in-degree, $\delta$ is a matrix of dyadic effects governing dyadic reciprocity, and the ellipse signifies any linear model of coefficients and focal, recipient, or dyadic covariates. For example, if $S$ is an animal-specific measure, like body size, and $Q$ is a dyad-specific measure, like a matrix of relatedness ties, then the ellipse may be replaced with: $\kappa_{[1]}S_{[i]} + \kappa_{[2]}S_{[j]} + \kappa_{[3]}Q_{[i,j]}$, to give the effects of body size on the probability of both sending and receiving transfers, and the effects of kinship on the probability of ties in either direction. In the Bernoulli models, slope parameters are indicative of changes in the the log-odds of tie probability, and block parameters are indicative of log-odds offsets. Researchers should be careful to evaluate the contrast between block-level parameters, to test for differences.

To complete the model definition, we model the sender and receiver effects jointly using a multivariate normal distribution (exactly as described in Redhead et al. 2021):

$$\begin{pmatrix} \lambda_{[i]} \\ \pi_{[i]} \end{pmatrix} \sim \text{MV Normal}\left( \begin{pmatrix} 0 \\ 0 \end{pmatrix}, \begin{pmatrix} \sigma_\lambda^2 & \sigma_\pi \sigma_\lambda \rho \\ \sigma_\lambda \sigma_\pi \rho & \sigma_\pi^2 \end{pmatrix} \right) \tag{8}$$

This allows for generalised correlations at the individual level to be detected—i.e., we can detect if individuals who groom others are also more likely to be groomed by others. For computational reasons (Stan Development Team 2021; Lewandowski et al. 2009), however, it is better to implement Eq. 8 by defining:

$$\begin{pmatrix} \lambda_{[i]} \\ \pi_{[i]} \end{pmatrix} = \begin{pmatrix} \sigma_\lambda \\ \sigma_\pi \end{pmatrix} \circ \left( L * \begin{pmatrix} \hat{\lambda}_{[i]} \\ \hat{\pi}_{[i]} \end{pmatrix} \right) \tag{9}$$

where $L$ is a Cholesky factor from the decomposition of the $2 \times 2$ correlation matrix with $\rho$ on the off-diagonal, and $\hat{\lambda}_{[i]} \sim \text{Normal}(0, 1)$ and $\hat{\pi}_{[i]} \sim \text{Normal}(0, 1)$ are unit-normal random effects. Weakly informative priors may then be independently specified on the variance and correlation terms (Lewandowski et al. 2009):

$$\sigma_\lambda \sim \text{Exponential}(1.5) \tag{10}$$
$$\sigma_\pi \sim \text{Exponential}(1.5) \tag{11}$$
$$L \sim \text{LKJ Cholesky}(2.0) \tag{12}$$

We use the above approach to define the dyad-level random effects as well:

$$\begin{pmatrix} \delta_{[i,j]} \\ \delta_{[j,i]} \end{pmatrix} = \begin{pmatrix} \sigma_\delta \\ \sigma_\delta \end{pmatrix} \circ \left( L_\delta * \begin{pmatrix} \hat{\delta}_{[i,j]} \\ \hat{\delta}_{[j,i]} \end{pmatrix} \right) \tag{13}$$

where $\hat{\delta}_{[i,j]} \sim \text{Normal}(0, 1)$ have unit-normal priors, and the variance and correlation terms have weakly informative priors:

$$\sigma_\delta \sim \text{Exponential}(1.5) \tag{14}$$
$$L_\delta \sim \text{LKJ Cholesky}(2.0) \tag{15}$$

Under this model, $\rho$ provides an indication of generalised reciprocity—i.e., whether those who give more (to others) also receive more (from others)—and $\rho_\delta$ provides a measure of dyadic reciprocity—i.e., whether the probability of focal $i$ giving to alter $j$, increases with the probability that focal $j$ gives to alter $i$.

##### 1.2. The Poisson model for counts

In many cases, researchers have data on the number of times ties specific kinds of ties were observed, but do not have an exposure variable. For example, researchers might count the number of times GPS trackers on a population of animals were within a 5-meter radius of each other over a standardised 1-month time period.

In this case, Eq. 6 is replaced by:

$$Y_{[i,j]} \sim \text{Poisson}\Big(\exp(\phi_{[i,j]})\Big) \tag{16}$$

while the linear model given in Eq. 7 remains unchanged:

$$\phi_{[i,j]} = \left(\sum_{v=0}^{V} B_{[v,b(i,v),b(j,v)]}\right) + \lambda_{[i]} + \pi_{[j]} + \delta_{[i,j]} + \dots \tag{17}$$

In the Poisson model, regression slopes are indicative of change in the log of expected counts as a function of change in the respective predictor variable.

##### 1.3. The Binomial model for proportions

In other cases, researchers do not simply have data on the presence or absence of ties, but instead have data on the number of times that ties *were observed* conditional on the number of times that ties *could have been observed*. For example, researchers may have conducted a specific number of scans, and the elements $Y_{[i,j]} \in \mathbb{N}$ might reflect the number of directed ties (e.g., resource transfers, or aggression events) from individual $i$ to individual $j$ that were observed. In this case, the number of scans is like an exposure variable, which limits the maximum possible count in the outcome. Thus, we let the variable $W_{[i,j]}$ be the count of scans in which individuals $i$ and $j$ could have engaged in the dyadic behaviour of interest.

In this case, Eq. 6 replaced by:

$$Y_{[i,j]} \sim \text{Binomial}\Big(W_{[i,j]}, \text{Logistic}(\phi_{[i,j]})\Big) \tag{18}$$

while the linear model given in Eq. 7 remains unchanged:

$$\phi_{[i,j]} = \left(\sum_{v=0}^{V} B_{[v,b(i,v),b(j,v)]}\right) + \lambda_{[i]} + \pi_{[j]} + \delta_{[i,j]} + \dots \tag{19}$$

As in the Bernoulli models, slope parameters in the Binomial models are indicative of changes in the log-odds of tie probability, and block parameters are indicative of log-odds offsets.

#### 2. Model validation

We conducted several simulation experiments to validate the performance of our models across a broad array of conditions, and to examine if there are any regions of parameter space where the models behave sub-optimally. To do this, we first generated network data using forward simulations from our stochastic blockmodel, our social relations model, and the combined model, which includes both stochastic blockmodel and social relations model parameters. We then used the corresponding inferential statistical models to analyse the simulated data and ensure that we can recover the generative parameter values. We conducted sweeps across all key model parameters, for each model, for each outcome mode (i.e., Bernoulli, Poisson, and Binomial). In each simulation experiment, we generally varied only a single generative parameter (e.g., the dyadic reciprocity rate) across a broad parameter space that contains realistic values, while fixing all other parameters in the model to empirically plausible values.

##### 2.1. Stochastic blockmodel validation

To begin our model validation procedure, we conduct a simulation experiment in which we generate data using the stochastic blockmodel simulation function provided in `STRAND` and then analyze these data using the corresponding model fitting and inference functions.

Figure 1 shows the test results for the Bernoulli model, Figure 2 shows the test results for the Poisson model, and Figure 3 shows the test results for the Binomial model. Predicted parameter values are plotted as yellow confidence regions, and the generative parameter values are shown in black.

Figure 1a, for example, shows the estimated intercepts for one within-block and one between-block tie as the model is fit to larger samples. We successfully recover the generative parameter values across a wide range of sample sizes; however, the precision of our estimates improves as sample size grows. We repeat this procedure in Figures 1b-1d, varying the generative parameter of interest (e.g., number of observed blocks, covariate effect size, etc.). Across all of these parameter sweeps, for each of the three outcome types, we accurately recover all generative parameter values.

[Fig. 1 about here.]

[Fig. 2 about here.]

[Fig. 3 about here.]

##### 2.2. Social relations model validation

To continue our model validation procedure, we conduct a simulation experiment in which we generate data using the social relations model simulation function provided in `STRAND` and then analyze these data using the corresponding model fitting and inference functions.

As with the stochastic blockmodel validation, we begin by demonstrating that our social relations model can accurately recover generative scalar parameters (as shown in Figures 4, 5, and 6, for the Bernoulli, Poisson, and Binomial outcomes, respectively).

[Fig. 4 about here.]

[Fig. 5 about here.]

[Fig. 6 about here.]

Then, because the social relations model includes individual- and dyad-level random effects, we also examine whether our model accurately recovers these parameters. To measure the similarity between the generative and estimated parameters, we calculate the correlation coefficient (Figures 7, 8, and 9). In general, the correlation plots suggest that we reliably recover focal random effects (governing out-degree) and target random effects (governing in-degree). Dyadic random effects are only recoverable in cases where the variance in dyadic random effects is high.

[Fig. 7 about here.]

[Fig. 8 about here.]

[Fig. 9 about here.]

##### 2.3. Combined social relations model and stochastic block model validation

The next step of our model validation procedure is to conduct a simulation experiment in which we generate data using the combined stochastic block model and social relations model simulation function provided in `STRAND` and then analyse these data using the corresponding model fitting and inference functions. As with the previous validations, we begin by demonstrating that this model can accurately recover generative scalar parameters (shown in Figures 10, 11, and 12).

[Fig. 10 about here.]

[Fig. 11 about here.]

[Fig. 12 about here.]

Then, because the combined model includes individual- and dyad-level random effects, we also examine whether our model accurately recovers these parameters. Specifically, we calculate the correlation coefficient between simulated and estimated random effects (Figures 13, 14, and 15). In general, the correlation plots suggest that we reliably recover focal offsets (governing out-degree), target offsets (governing in-degree), and even dyadic offsets (because of block structure).

[Fig. 13 about here.]

[Fig. 14 about here.]

[Fig. 15 about here.]

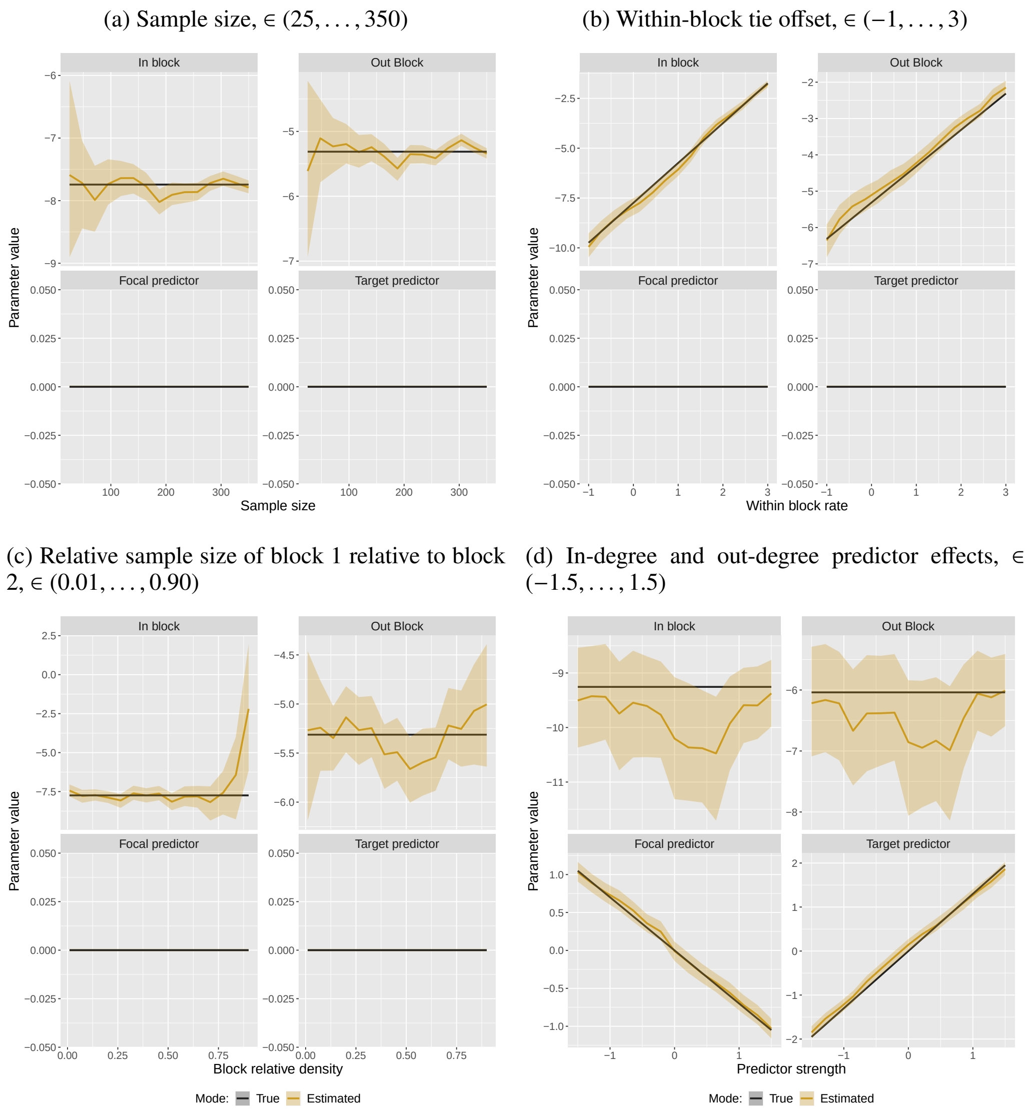

(a) Sample size, $\in (25, \ldots, 350)$

(b) Within-block tie offset, $\in (-1, \ldots, 3)$

(c) Relative sample size of block 1 relative to block 2, $\in (0.01, \ldots, 0.90)$

(d) In-degree and out-degree predictor effects, $\in (-1.5, \ldots, 1.5)$

Fig. 1: Parameter recovery from the stochastic block model, Bernoulli outcomes. Each frame plots the parameter values estimated by our model (in yellow), and the true values specified in the simulation (in black). The y-axis of each sub-figure represents the parameter value, and the x-axis represents the value of the focal simulation parameter given in the sub-heading. For example, the x-axis in Figure 1a represents the sample size used when simulating the network data.

(a) Sample size, $\in (25, \ldots, 350)$

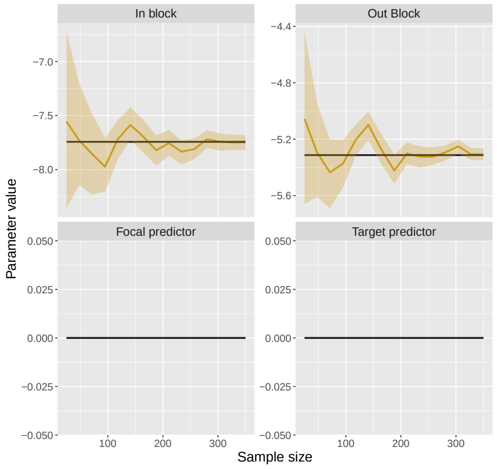

(b) Within-block tie offset, $\in (-1, \ldots, 3)$

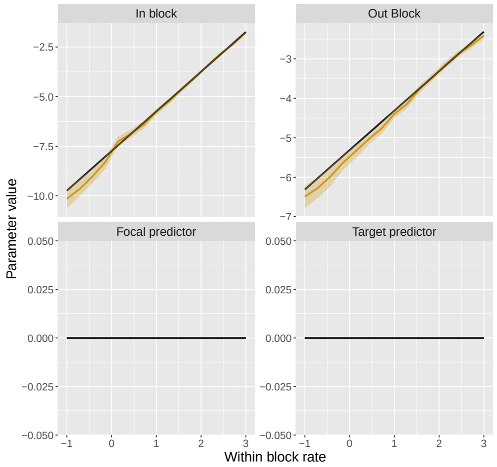

(c) Relative sample size of block 1 relative to block 2, $\in (0.01, \ldots, 0.90)$

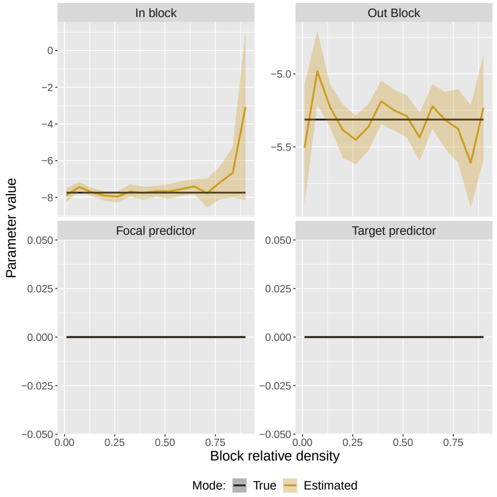

(d) In-degree and out-degree predictor effects, $\in (-1.5, \ldots, 1.5)$

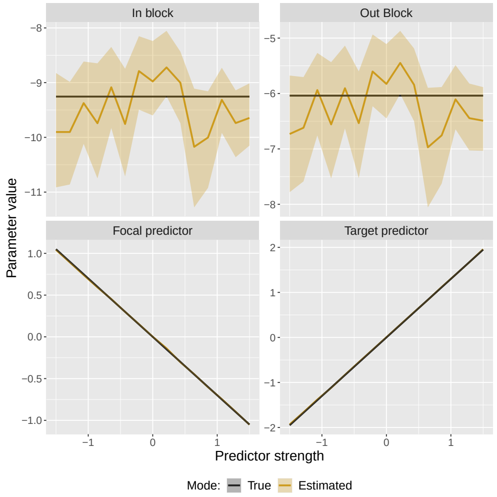

Fig. 2: Parameter recovery from the stochastic block model, Poisson outcomes. Each frame plots the parameter values estimated by our model (in yellow), and the true values specified in the simulation (in black). The y-axis of each sub-figure represents the parameter value, and the x-axis represents the value of the focal simulation parameter given in the sub-heading. For example, the x-axis in Figure 2a represents the sample size used when simulating the network data.

(a) Sample size, $\in (25, \ldots, 350)$

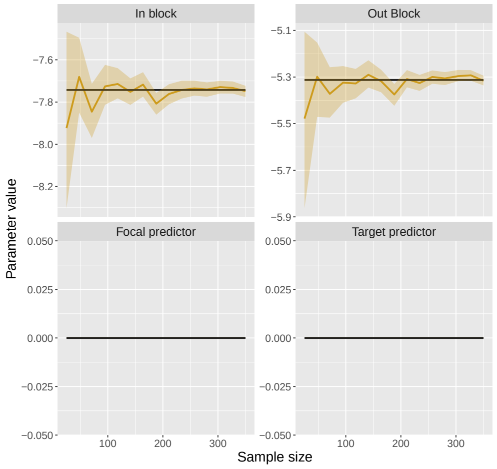

(b) Within-block tie offset, $\in (-1, \ldots, 3)$

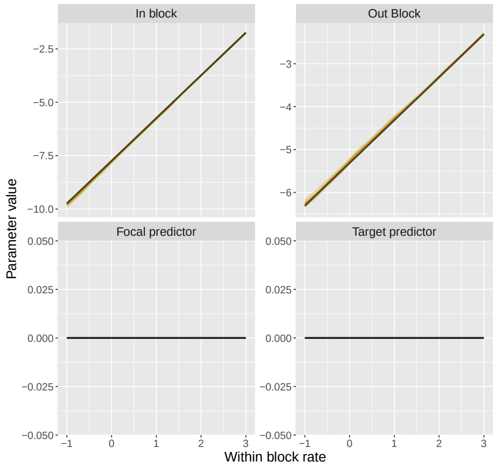

(c) Relative sample size of block 1 relative to block 2, $\in (0.01, \ldots, 0.90)$

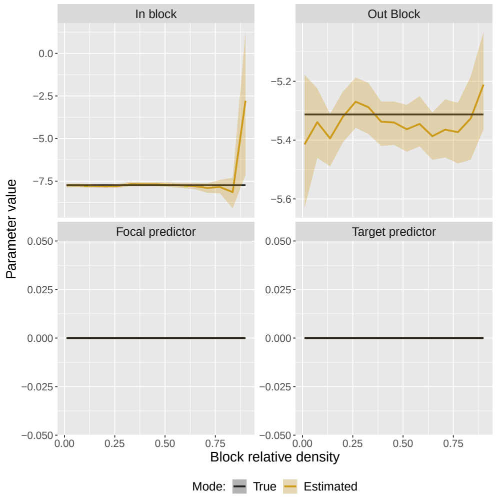

(d) In-degree and out-degree predictor effects, $\in (-1.5, \ldots, 1.5)$

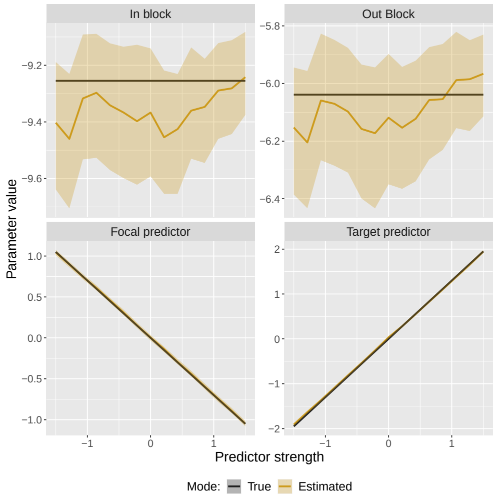

Fig. 3: Parameter recovery from the stochastic block model, Binomial outcomes. Each frame plots the parameter values estimated by our model (in yellow), and the true values specified in the simulation (in black). The y-axis of each sub-figure represents the parameter value, and the x-axis represents the value of the focal simulation parameter given in the sub-heading. For example, the x-axis in Figure 3a represents the sample size used when simulating the network data.

(a) Intercept log-odds, ∈ (−10, . . . , 2)

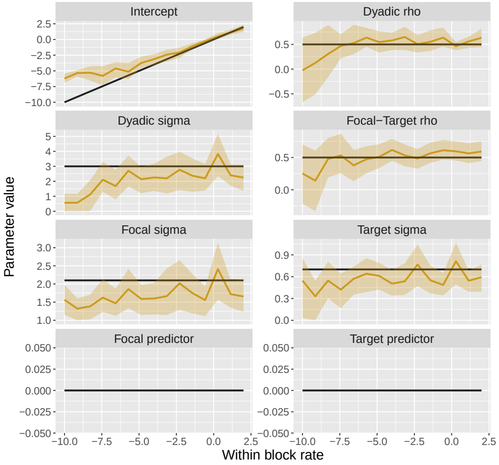

(b) In- and out-degree predictor effects, ∈ (−2, . . . , 2)

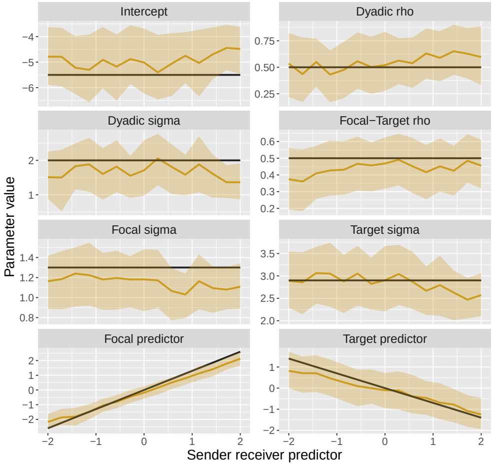

(c) Correlation of in- and out-degree (generalized reciprocity), ∈ (−0.8, . . . , 0.8)

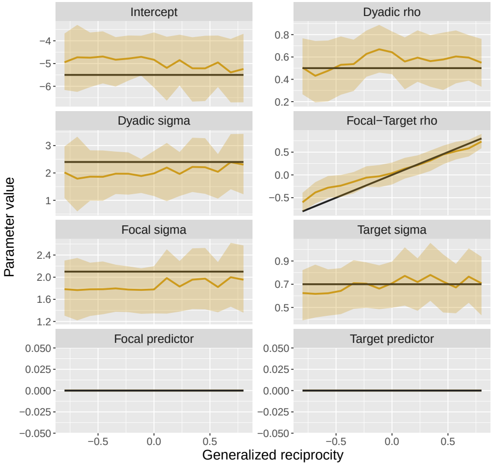

(d) SD of in- and out-degree, ∈ (0.28, . . . , 3.8)

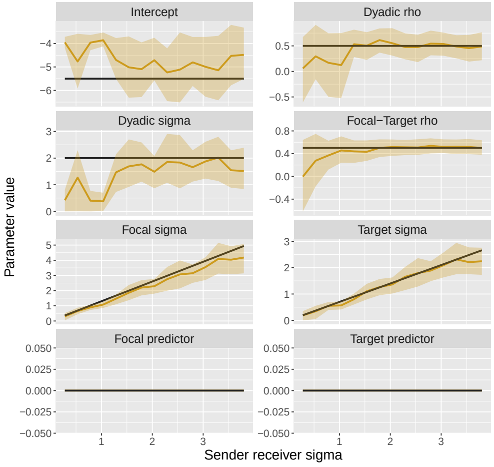

(e) Correlation of dyadic effects (dyadic reciprocity), ∈ (0.01, . . . , 0.80)

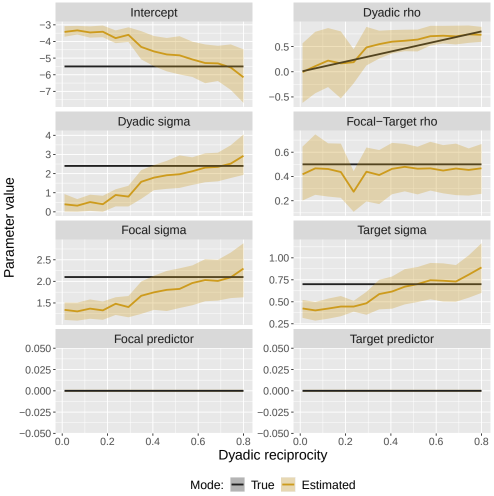

(f) SD of dyadic effects, ∈ (0.28, . . . , 3.8)

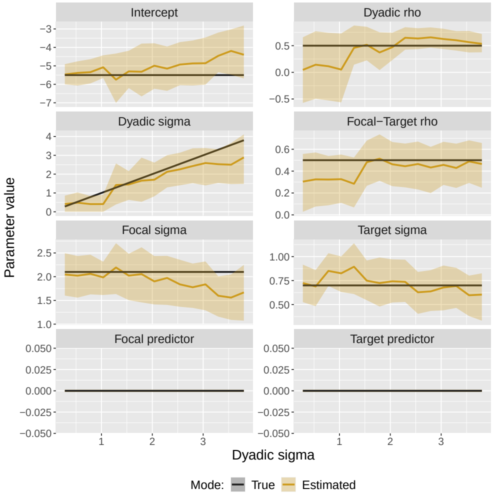

Fig. 4: Parameter recovery from the social relations model, Bernoulli outcomes. Each frame plots the parameter values estimated by our model (in yellow), and the true values specified in the simulation (in black). The y-axis of each sub-figure represents the parameter value, and the x-axis represents the value of the focal simulation parameter given in the sub-heading. For example, the x-axis in Figure 4a represents the intercept log-odds used when simulating the network data.

(a) Intercept log-odds, ∈ $(-10, \ldots, 2)$

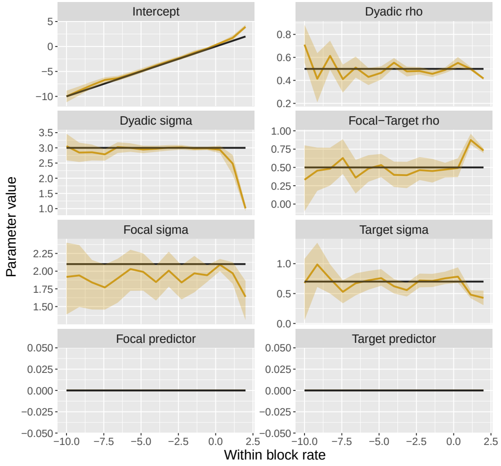

(b) In- and out-degree predictor effects, ∈ $(-2, \ldots, 2)$

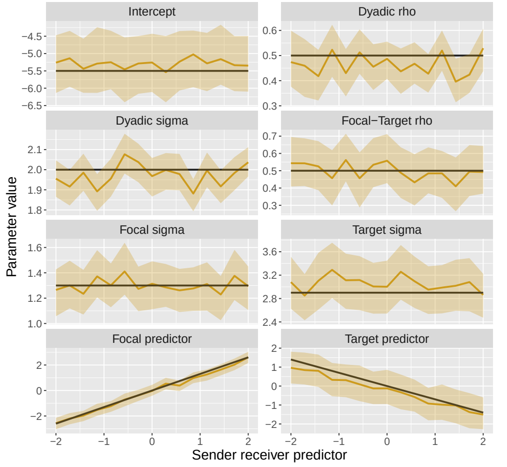

(c) Correlation of in- and out-degree (generalized reciprocity), ∈ $(-0.8, \ldots, 0.8)$

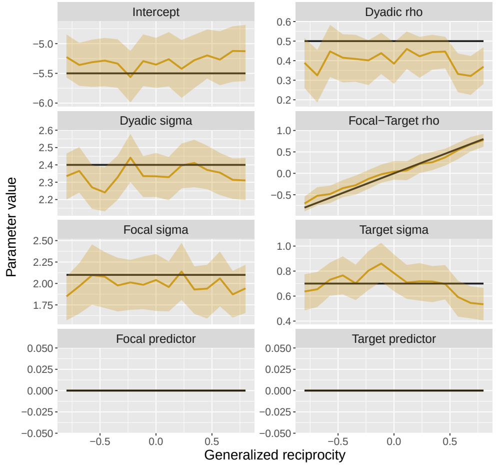

(d) SD of in- and out-degree, ∈ $(0.28, \ldots, 3.8)$

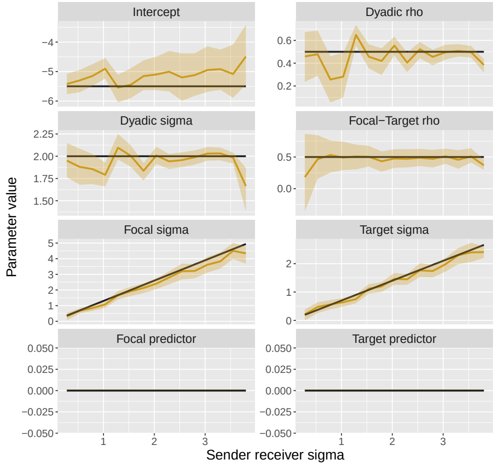

(e) Correlation of dyadic effects (dyadic reciprocity), ∈ $(0.01, \ldots, 0.80)$

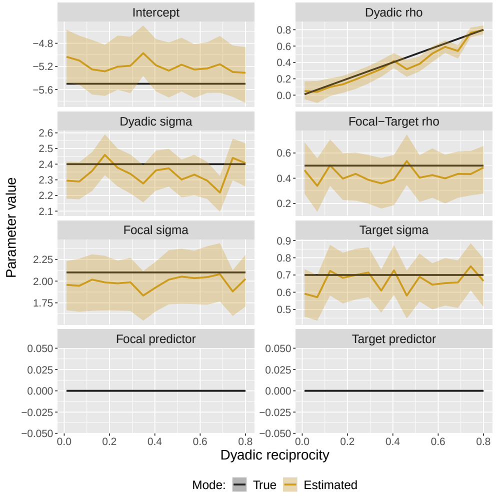

(f) SD of dyadic effects, ∈ $(0.28, \ldots, 3.8)$

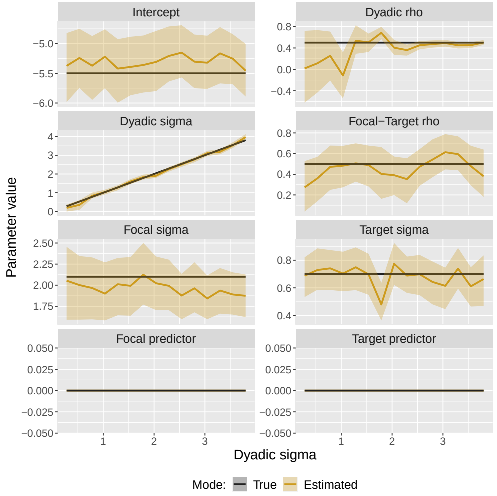

Fig. 5: Parameter recovery from the social relations model, Poisson outcomes. Each frame plots the parameter values estimated by our model (in yellow), and the true values specified in the simulation (in black). The y-axis of each sub-figure represents the parameter value, and the x-axis represents the value of the focal simulation parameter given in the sub-heading. For example, the x-axis in Figure 5a represents the intercept log-odds used when simulating the network data.

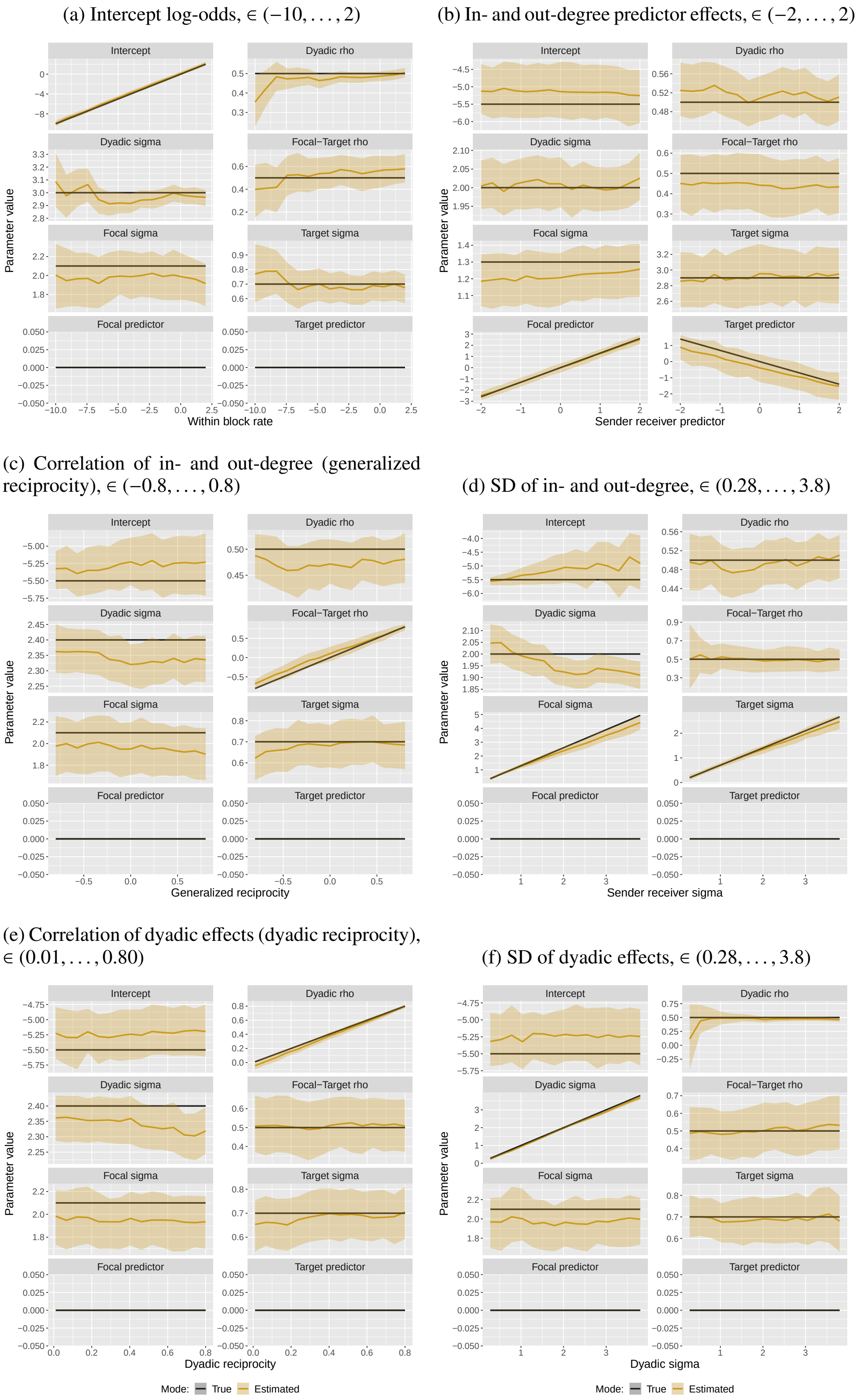

Fig. 6: Parameter recovery from the social relations model, Binomial outcomes. Each frame plots the parameter values estimated by our model (in yellow), and the true values specified in the simulation (in black). The y-axis of each sub-figure represents the parameter value, and the x-axis represents the value of the focal simulation parameter given in the sub-heading. For example, the x-axis in Figure 6a represents the intercept log-odds used when simulating the network data.

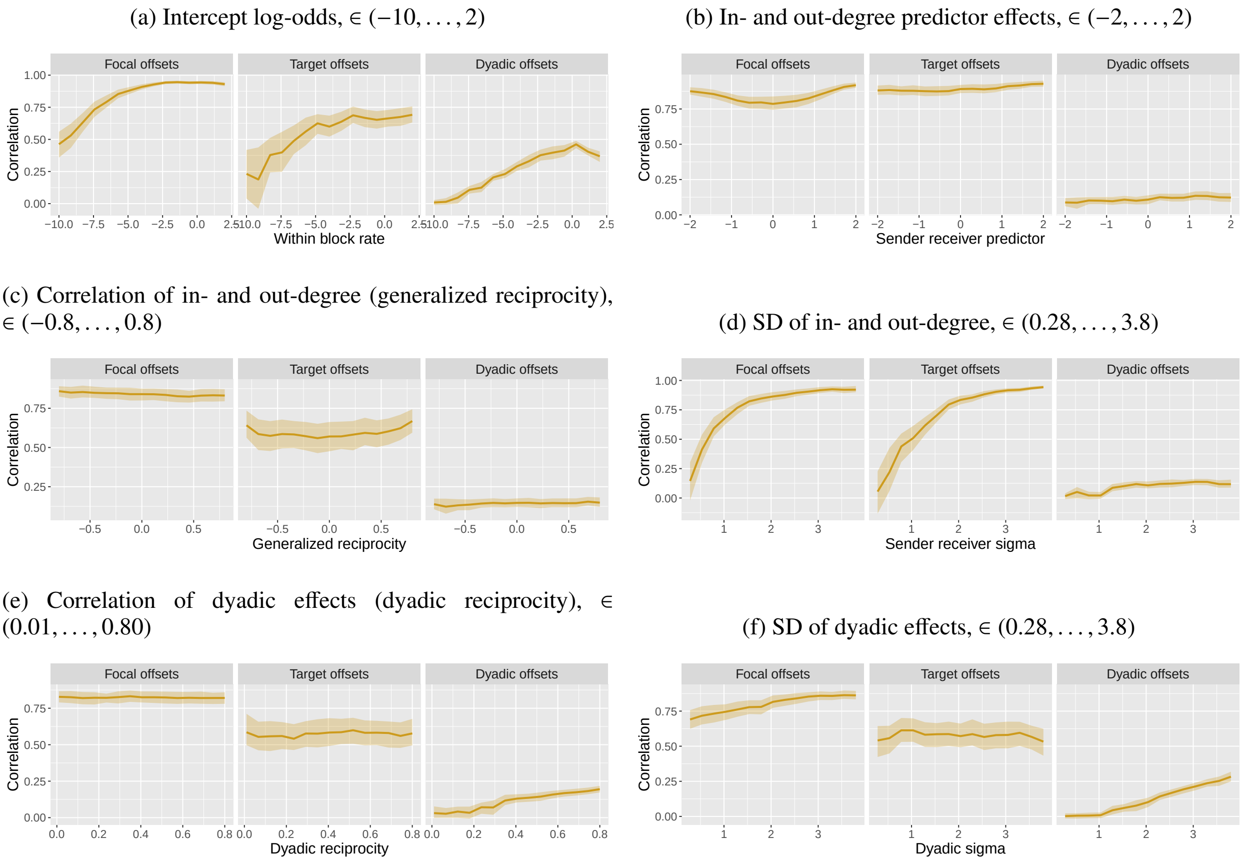

Fig. 7: Parameter recovery from the social relations model, Bernoulli outcomes. Each frame plots the correlation (in yellow) between the generative random effects and the estimated random effects. The y-axis of each sub-figure represents the correlation value, and the x-axis represents the value of the focal simulation parameter given in the sub-heading.

(a) Intercept log-odds, $\in (-10, \ldots, 2)$

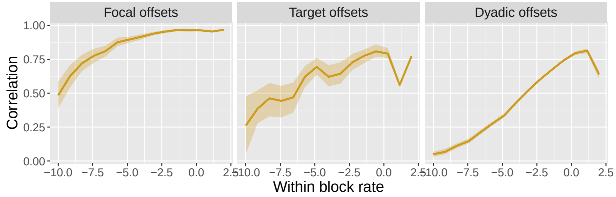

(b) In- and out-degree predictor effects, $\in (-2, \ldots, 2)$

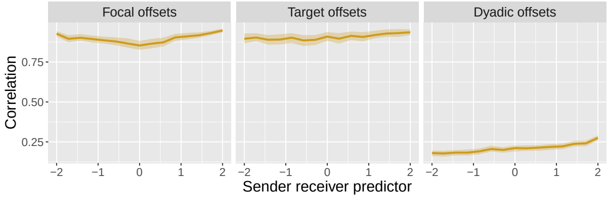

(c) Correlation of in- and out-degree (generalized reciprocity), $\in (-0.8, \ldots, 0.8)$

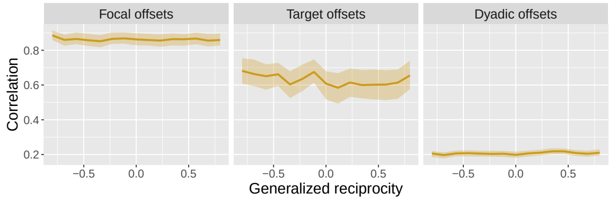

(d) SD of in- and out-degree, $\in (0.28, \ldots, 3.8)$

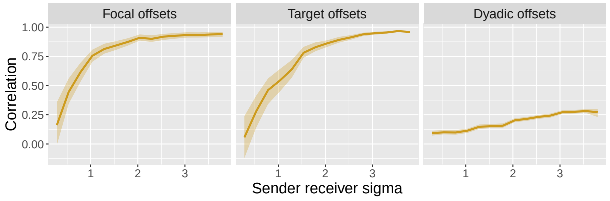

(e) Correlation of dyadic effects (dyadic reciprocity), $\in (0.01, \ldots, 0.80)$

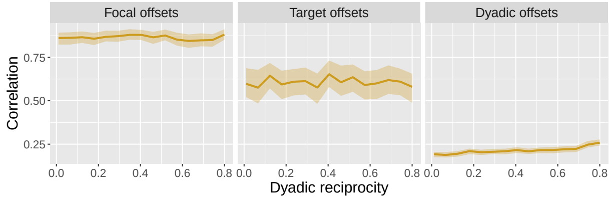

(f) SD of dyadic effects, $\in (0.28, \ldots, 3.8)$

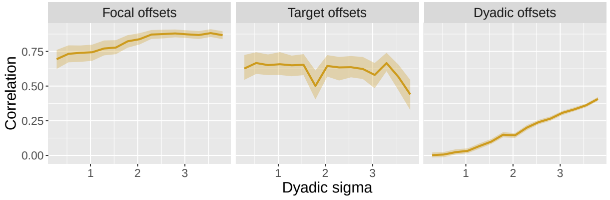

Fig. 8: Parameter recovery from the social relations model, Poisson outcomes. Each frame plots the correlation (in yellow) between the generative random effects and the estimated random effects. The y-axis of each sub-figure represents the correlation value, and the x-axis represents the value of the focal simulation parameter given in the sub-heading.

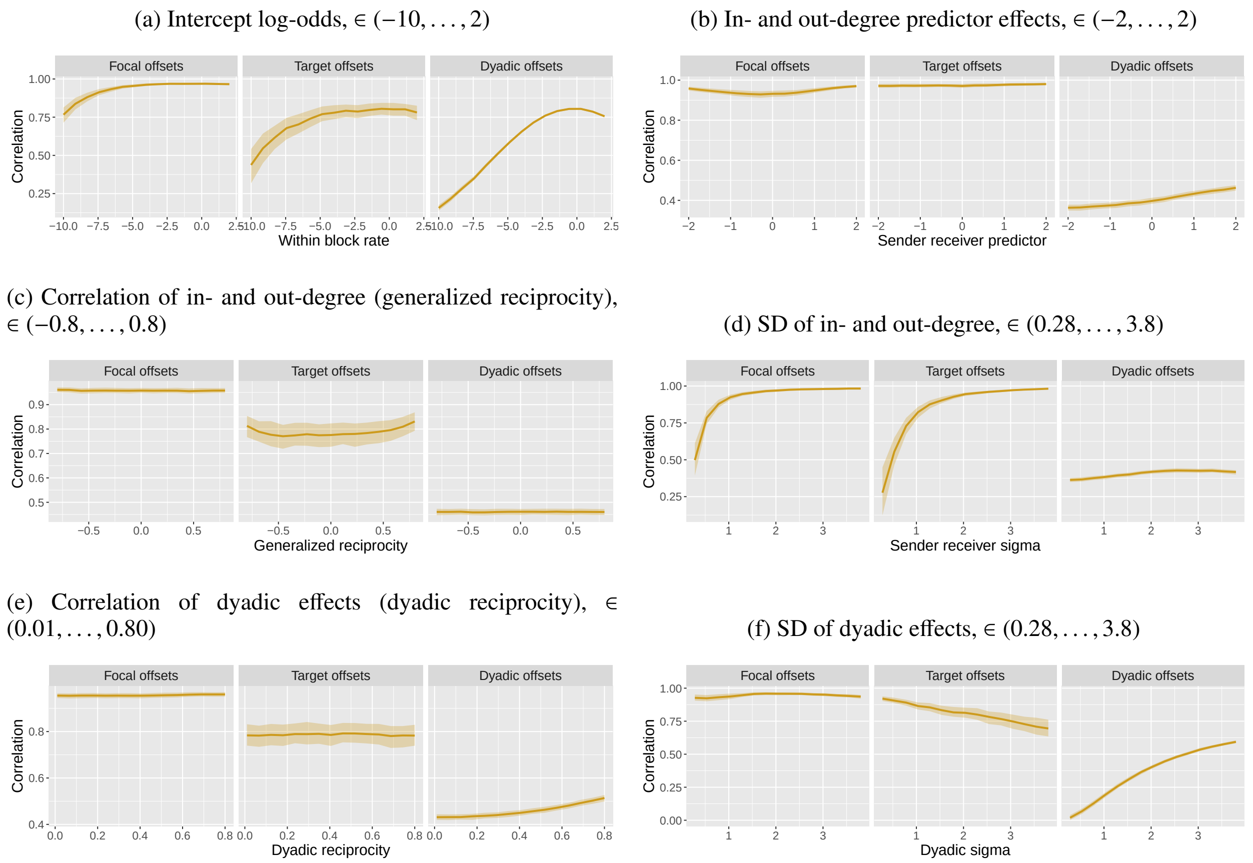

Fig. 9: Parameter recovery from the social relations model, Binomial outcomes. Each frame plots the correlation (in yellow) between the generative random effects and the estimated random effects. The y-axis of each sub-figure represents the correlation value, and the x-axis represents the value of the focal simulation parameter given in the sub-heading.

(a) Intercept offset log-odds, ∈ (−4, . . . , 6)

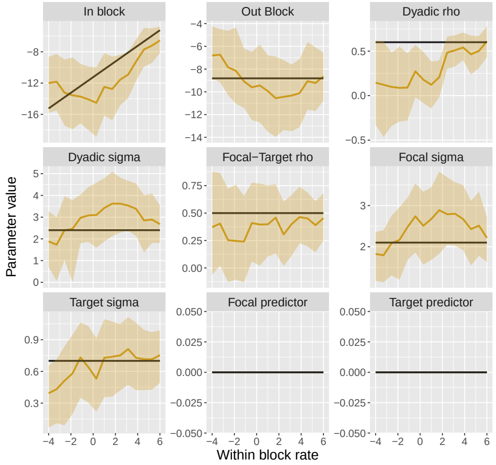

(b) In- and out-degree predictor effects, ∈ (−2, . . . , 2)

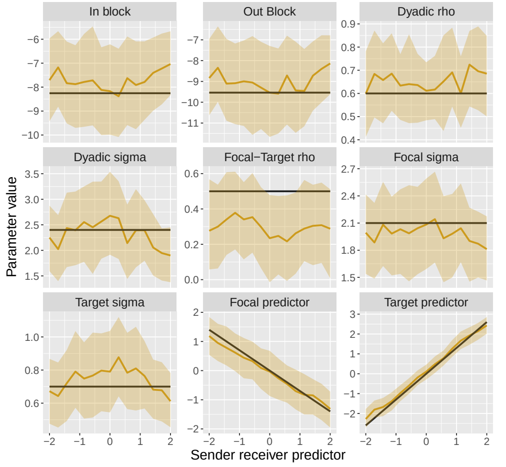

(c) Correlation of in- and out-degree (generalized reciprocity), ∈ (−0.8, . . . , 0.8)

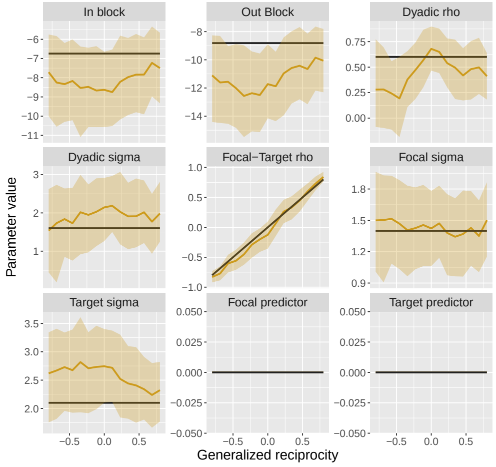

(d) SD of in- and out-degree, ∈ (0.3, . . . , 3.8)

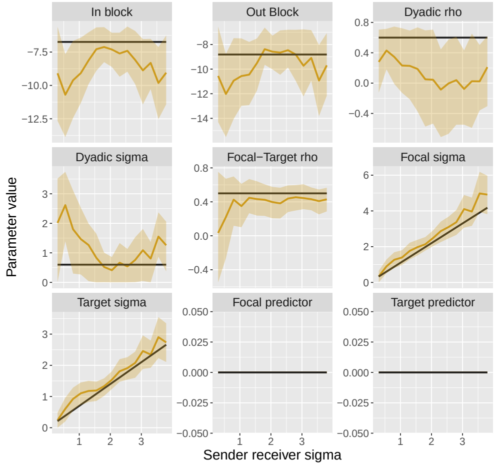

(e) Correlation of dyadic effects (dyadic reciprocity), ∈ (0.01, . . . , 0.80)

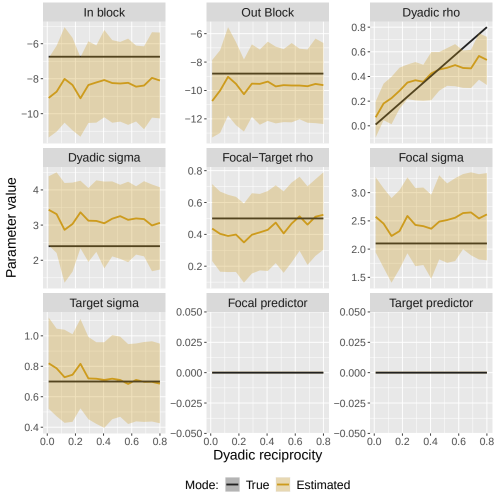

(f) SD of dyadic effects, ∈ (0.3, . . . , 3.8)

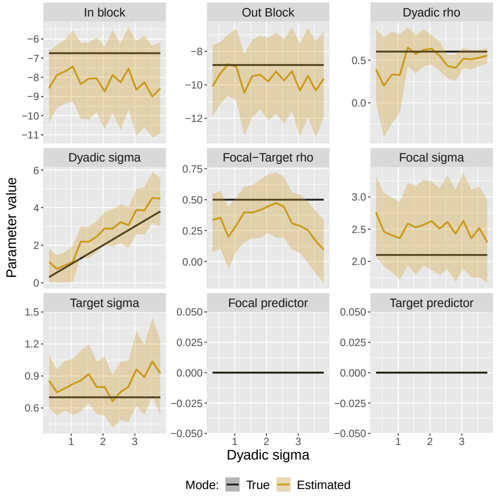

Fig. 10: Parameter recovery from the joint block and social relations model, Bernoulli outcomes. Each frame plots the parameter values estimated by our model (in yellow), and the true values specified in the simulation (in black). The y-axis of each sub-figure represents the parameter value, and the x-axis represents the value of the focal simulation parameter given in the sub-heading. For example, the x-axis in Figure 10a represents the within-block intercept offset in log-odds used when simulating the network data.

(a) Intercept offset log-odds, $\in (-4, \ldots, 6)$

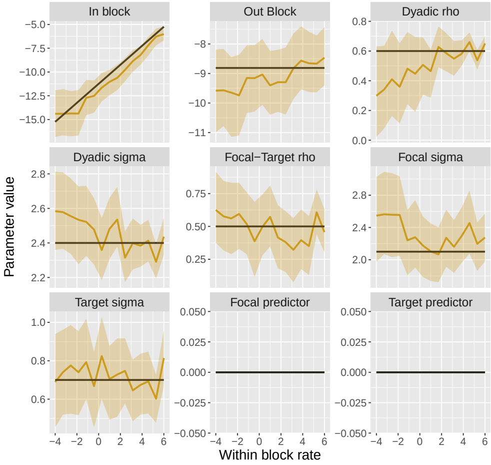

(b) In- and out-degree predictor effects, $\in (-2, \ldots, 2)$

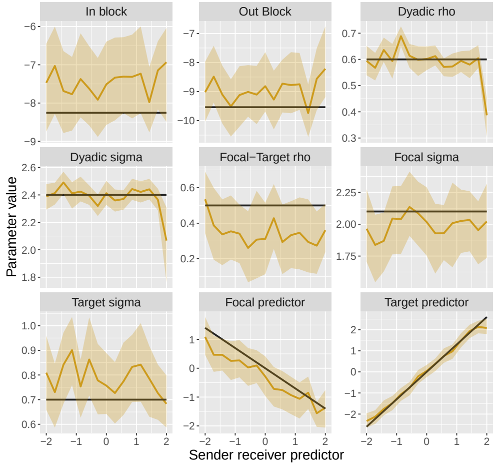

(c) Correlation of in- and out-degree (generalized reciprocity), $\in (-0.8, \ldots, 0.8)$

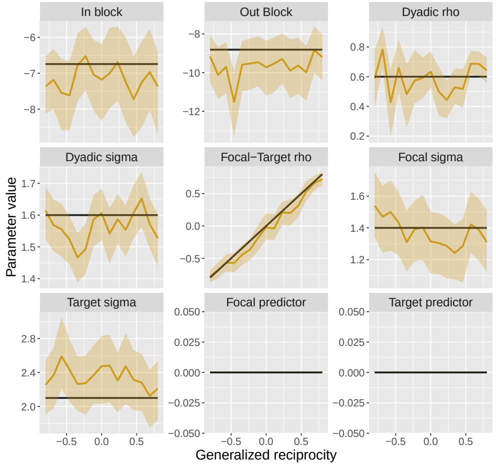

(d) SD of in- and out-degree, $\in (0.3, \ldots, 3.8)$

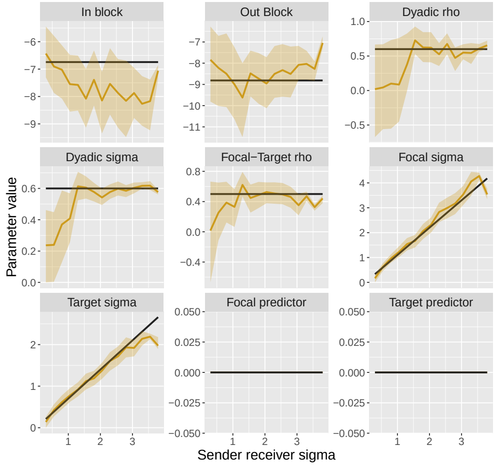

(e) Correlation of dyadic effects (dyadic reciprocity), $\in (0.01, \ldots, 0.80)$

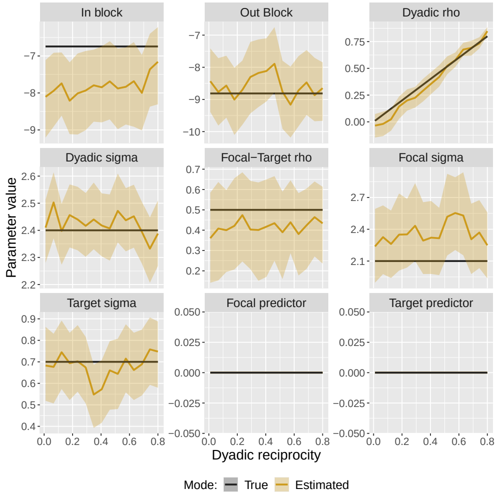

(f) SD of dyadic effects, $\in (0.3, \ldots, 3.8)$

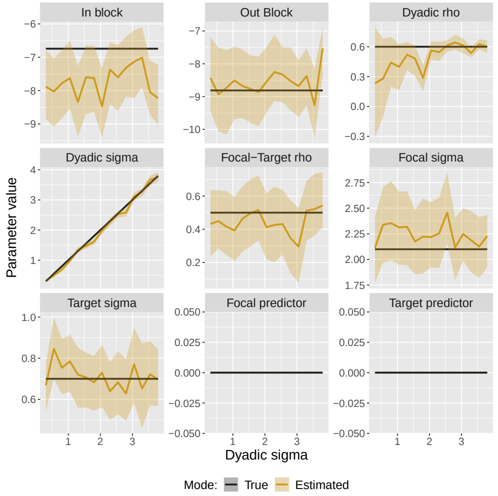

Fig. 11: Parameter recovery from the joint block and social relations model, Poisson outcomes. Each frame plots the parameter values estimated by our model (in yellow), and the true values specified in the simulation (in black). The y-axis of each sub-figure represents the parameter value, and the x-axis represents the value of the focal simulation parameter given in the sub-heading. For example, the x-axis in Figure 11a represents the within-block intercept offset in log-odds used when simulating the network data.

(a) Intercept offset log-odds, $\in (-4, \ldots, 6)$

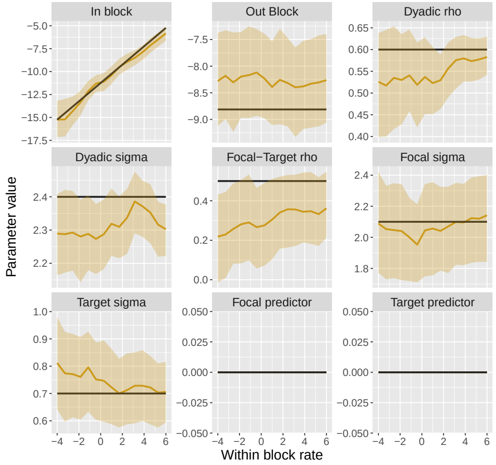

(b) In- and out-degree predictor effects, $\in (-2, \ldots, 2)$

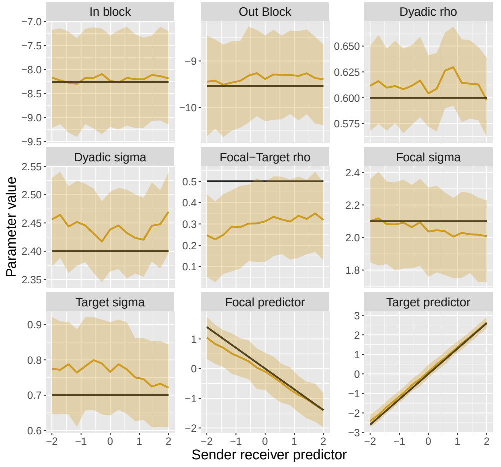

(c) Correlation of in- and out-degree (generalized reciprocity), $\in (-0.8, \ldots, 0.8)$

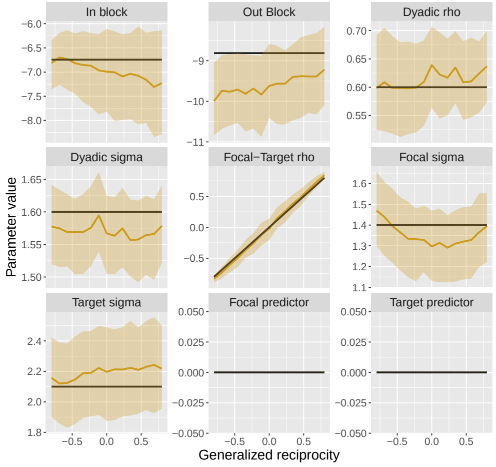

(d) SD of in- and out-degree, $\in (0.3, \ldots, 3.8)$

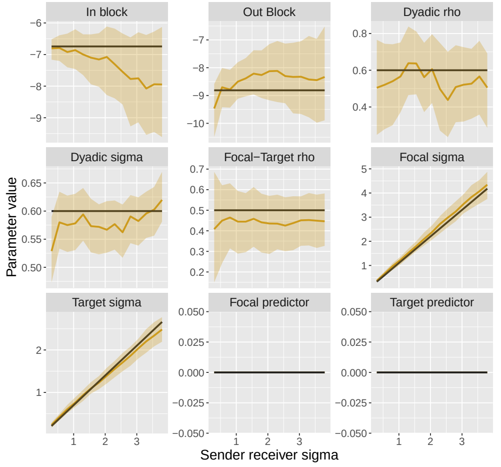

(e) Correlation of dyadic effects (dyadic reciprocity), $\in (0.01, \ldots, 0.80)$

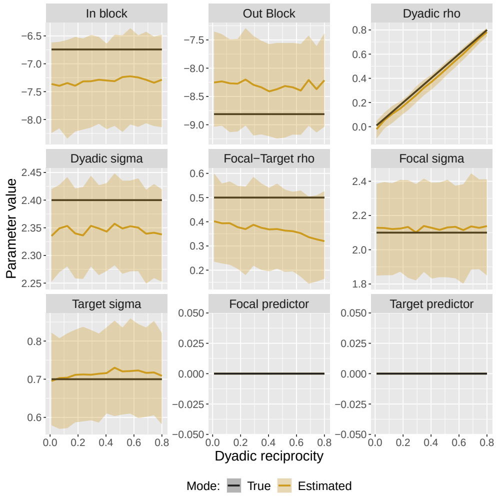

(f) SD of dyadic effects, $\in (0.3, \ldots, 3.8)$

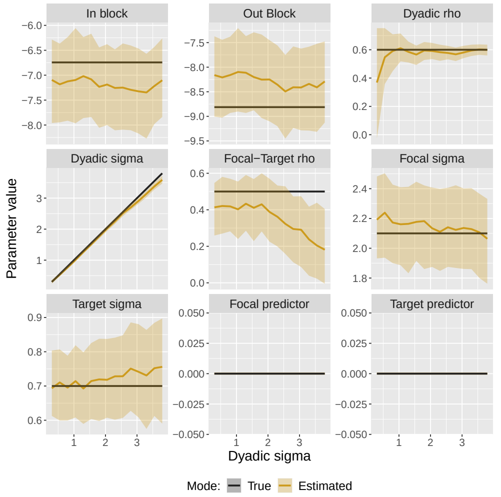

Fig. 12: Parameter recovery from the joint block and social relations model, Binomial outcomes. Each frame plots the parameter values estimated by our model (in yellow), and the true values specified in the simulation (in black). The y-axis of each sub-figure represents the parameter value, and the x-axis represents the value of the focal simulation parameter given in the sub-heading. For example, the x-axis in Figure 12a represents the within-block intercept offset in log-odds used when simulating the network data.

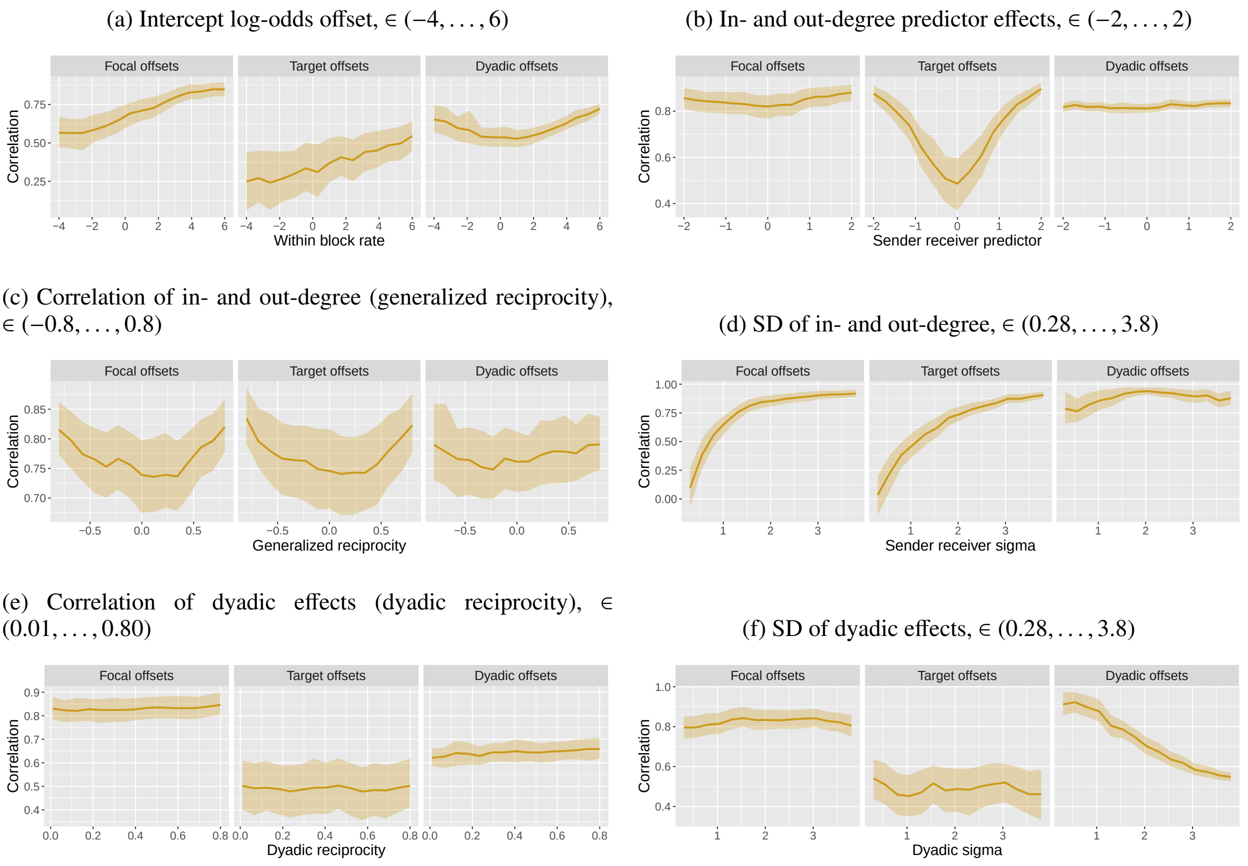

Fig. 13: Parameter recovery from the joint block and social relations model, Bernoulli outcomes. Each frame plots the correlation (in yellow) between the generative random effects and the estimated random effects. The y-axis of each sub-figure represents the correlation value, and the x-axis represents the value of the focal simulation parameter given in the sub-heading.

(a) Intercept log-odds offset, $\in (-4, \ldots, 6)$

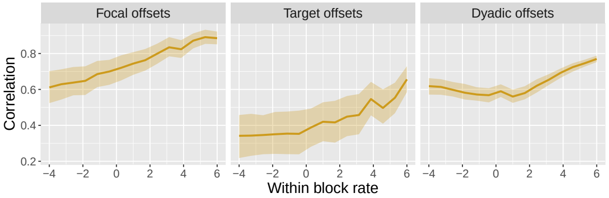

(b) In- and out-degree predictor effects, $\in (-2, \ldots, 2)$

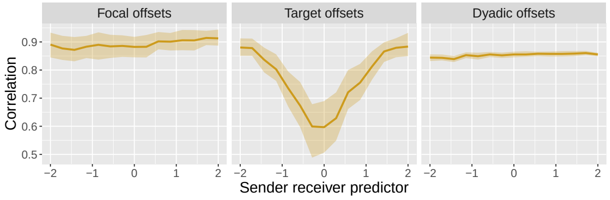

(c) Correlation of in- and out-degree (generalized reciprocity), $\in (-0.8, \ldots, 0.8)$

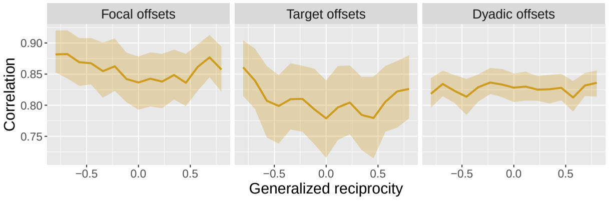

(d) SD of in- and out-degree, $\in (0.28, \ldots, 3.8)$

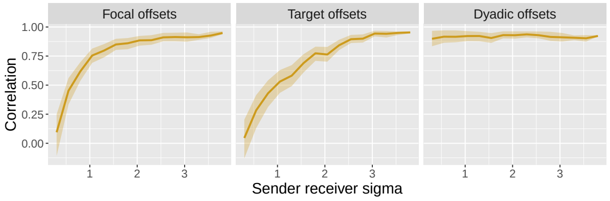

(e) Correlation of dyadic effects (dyadic reciprocity), $\in (0.01, \ldots, 0.80)$

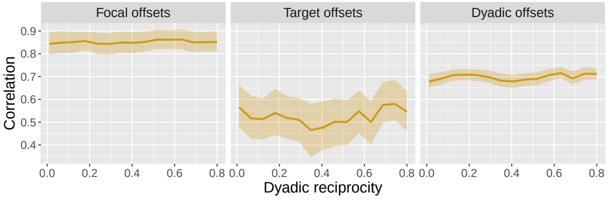

(f) SD of dyadic effects, $\in (0.28, \ldots, 3.8)$

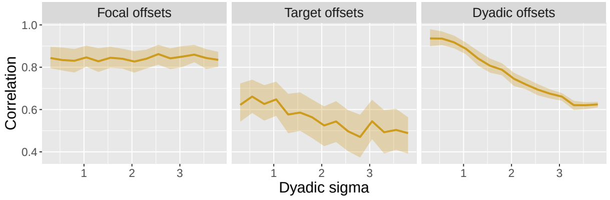

Fig. 14: Parameter recovery from the joint block and social relations model, Poisson outcomes. Each frame plots the correlation (in yellow) between the generative random effects and the estimated random effects. The y-axis of each sub-figure represents the correlation value, and the x-axis represents the value of the focal simulation parameter given in the sub-heading.

(a) Intercept log-odds offset, $\in (-4, \ldots, 6)$

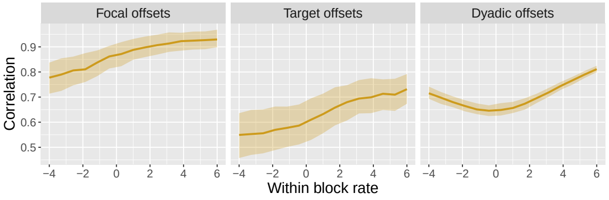

(b) In- and out-degree predictor effects, $\in (-2, \ldots, 2)$

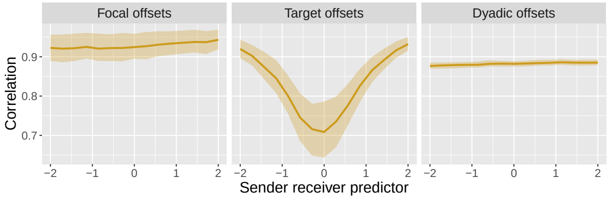

(c) Correlation of in- and out-degree (generalized reciprocity), $\in (-0.8, \ldots, 0.8)$

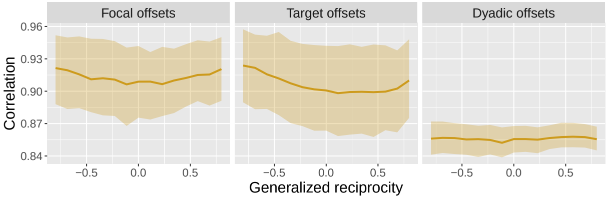

(d) SD of in- and out-degree, $\in (0.28, \ldots, 3.8)$

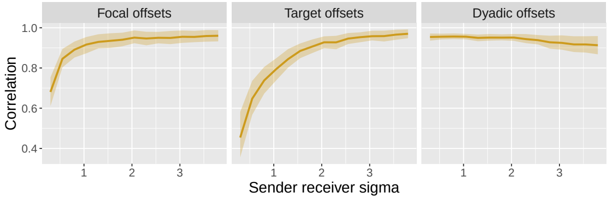

(e) Correlation of dyadic effects (dyadic reciprocity), $\in (0.01, \ldots, 0.80)$

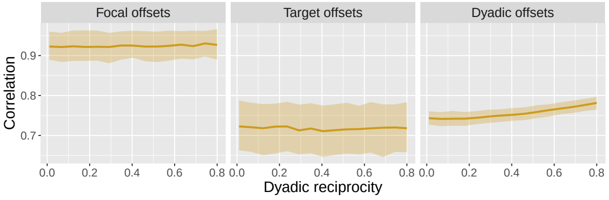

(f) SD of dyadic effects, $\in (0.28, \ldots, 3.8)$

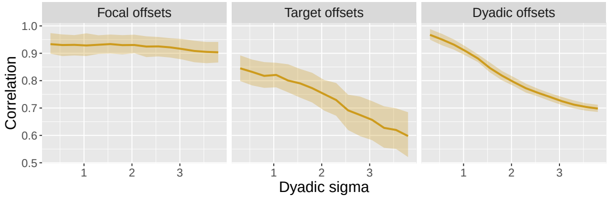

Fig. 15: Parameter recovery from the joint block and social relations model, Binomial outcomes. Each frame plots the correlation (in yellow) between the generative random effects and the estimated random effects. The y-axis of each sub-figure represents the correlation value, and the x-axis represents the value of the focal simulation parameter given in the sub-heading.
